# Seasonal changes in membrane structure and excitability in central neurons of goldfish (*Carassius auratus*) under constant environmental conditions

**DOI:** 10.1101/2022.03.04.483033

**Authors:** Michael W. Country, Kristina Haase, Katrin Blank, Carlos R. Canez, Joshua A. Roberts, Benjamin F.N. Campbell, Jeffrey C. Smith, Andrew E. Pelling, Michael G. Jonz

## Abstract

Seasonal modifications in the structure of cellular membranes occur as an adaptive measure to withstand exposure to prolonged environmental change. Little is known about whether such changes may occur independently of external cues, such as photoperiod or temperature, or how they may impact the central nervous system (CNS). We compared membrane properties of central neurons isolated from the retina of goldfish (*Carassius auratus*), an organism well-adapted to extreme environmental change, during the summer and winter months. Goldfish were maintained in a facility under constant environmental conditions throughout the year. Analysis of whole-retina phospholipid composition using mass spectrometry-based lipidomics revealed a two-fold increase in phosphatidylethanolamine species during the winter, suggesting an increase in cell membrane fluidity. Atomic force microscopy was used to produce localized, nanoscale-force deformation of neuronal membranes. Measurement of Young’s modulus indicated increased membrane stiffness (or decreased elasticity) in neurons isolated during the winter. Voltage-clamp electrophysiology was used to assess physiological changes in neurons between seasons. Winter neurons displayed a hyperpolarized reversal potential (*V*_rev_) and a significantly lower input resistance (*R*_in_) compared to summer neurons. This was indicative of a decrease in membrane excitability during the winter. Subsequent measurement of intracellular Ca^2+^ activity using Fura-2 microspectrofluorometry confirmed a reduction in action potential activity, including duration and action potential profile, in neurons isolated during the winter. These studies demonstrate chemical and biophysical changes that occur in central neurons of goldfish throughout the year without exposure to seasonal cues, and suggest a novel mechanism of seasonal regulation of CNS activity.

**SUMMARY STATEMENT:** Central neurons isolated from the retina of goldfish held under constant environmental conditions undergo seasonal changes in membrane structure and excitability.

## INTRODUCTION

Many organisms have evolved adaptations that allow them to survive extreme changes in in their environment. In the central nervous system (CNS) of anoxia-tolerant vertebrates, the matching of ATP supply to demand and the maintenance of ionic gradients across the plasma membrane occur even when oxygen is limiting (Bickler and Buck, 2007). Some species of fresh-water turtle (*Chrysemys picta* and *Trachemys scripta*), frogs (*Rana* spp.), the goldfish (*Carassius auratus*) and crucian carp (*C. carassius*) are remarkably tolerant to long periods of anoxia, especially at low temperatures (Herbert and Jackson, 1985; Jackson et al., 1984; van den Thillart et al., 1980; Lutz and Nilsson, 2004). Cellular homeostasis under such conditions may be accomplished by large-scale reductions in metabolic rate (Hochachka et al., 1996), increased tissue storage of fermentable substrate (Jackson and Ultsch, 2010) or ion channel arrest (Pamenter et al., 2008; Zivkovic and Buck, 2010; Rodgers-Garlick et al., 2013). These adaptations help to prevent the onset of anoxic injury to cells and tissues.

Long-term changes in temperature can induce modification in the phospholipid composition of the plasma membrane through homeoviscous adaptation (HVA), where a relative increase in saturated fatty acid composition is associated with an increase in environmental temperature (Hazel, 1995; Ernst et al., 2016). HVA was first described in bacteria (Sinensky, 1974) and later in cells derived from animals, including the goldfish brain (Cossins et al., 1981; Hazel and Williams, 1990). It is particularly important in poikilothermic organisms, as it leads to the synthesis of phospholipids to maintain constant viscosity or fluidity within the membrane as temperatures change. Phosphatidylethanolamines (PEs) are a structural component of membrane bilayers in all eukaryotes and comprise approximately 45% of total membrane phospholipids in the brain (Vance and Tasseva, 2013). PEs are particularly important in regulating membrane fluidity (Hazel, 1988; Pruitt, 1988; Hazel, 1995). Importantly, changes in membrane structure may have profound effects upon integral membrane protein activity. In vertebrates, the degree of polyunsaturation in membrane phospholipids is positively correlated with Na^+^/K^+^-ATPase activity and maintenance of ionic gradients (Cossins et al., 1981; Hulbert and Else, 1999, 2005). Moreover, changes in lipid bilayer composition, thickness, viscosity and curvature can affect the activity of other membrane proteins, including ion channels and G-protein-coupled receptors (Lee, 2004; Andersen and Koeppe, 2007; Rusinova et al., 2014; Fried et al., 2021).

The above adaptive chemical and physiological changes typically occur as a response to environmental cues, such as changes in temperature or oxygen availability that may occur during acclimatization or seasonal transitions. Interestingly, no studies have investigated whether seasonal changes at the level of the plasma membrane might occur independently of environmental cues, and what impact such changes may have upon the activity of central neurons. Based on the authors’ preliminary observations that the electrical properties of neurons isolated from goldfish kept in our animal facility differed throughout the calendar year, we designed a study carried out over multiple years to test the hypothesis that neurons isolated from animals held under constant environmental conditions during the summer versus winter months would display membranes with different biochemical and biophysical properties. We isolated neurons from the goldfish retina, a part of the CNS, and used mass spectrometry, atomic force microscopy, electrophysiology and microspectrofluorometry to probe the structural and functional characteristics in neuronal membranes. We discovered significant changes in membrane phospholipid composition, membrane stiffness, electrical excitability and action potential profile. These results indicate significant seasonal modifications in neuronal membrane structure and function in the absence of seasonal cues in an animal that has adapted to survive extreme environmental change.

## MATERIALS AND METHODS

### Animals

Adult common goldfish (*Carassius auratus*, Linnaeus 1758) of mixed age and sex weighing 10–75 g were used in the present study. Animals were obtained from commercial suppliers (Mirdo Importations Canada, Montréal, QC, Canada; or Aquality Tropical Fish Supply Inc., Mississauga, ON, Canada). Goldfish were originally sourced from sites in the southern United States, where they were bred and raised in outdoor ponds on a natural photoperiod. Upon arrival, animals were quarantined for 6 weeks and housed for up to 18 months in the aquatic facilities of the Laboratory for the Physiology and Genetics of Aquatic Organisms at the University of Ottawa, where they were maintained in 170 l tanks fitted with a flow-through system of fresh, aerated and dechloraminated water at a constant temperature of 18°C. For electrophysiological experiments, goldfish were obtained from the Aquatron Laboratory (Dalhousie University, Halifax, NS, Canada) and placed in a 400 l tank at 15°C. In both facilities, photoperiod was kept at a constant cycle of 12 h light:12 h dark and all environmental conditions were kept constant year-round. Before dissection and cell isolation, goldfish were dark adapted for approximately 1 h to facilitate removal of the retina, euthanized by rapid decapitation and pithed. Procedures for animal care and use were approved by the University of Ottawa Animal Care and Veterinary Services, and by the University Committee for Laboratory Animals at Dalhousie University. All procedures were implemented in accordance with regulations of the Canadian Council on Animal Care.

### Isolated cell preparation

Horizontal cells were isolated from goldfish retina following the procedures of Jonz and Barnes (2007). The retina is a convenient source of central neurons (Dowling, 1987), and horizontal cells are neurons whose membrane and physiological properties are well characterized (e.g. Tachibana, 1981; Lasater et al., 1984; Cunningham and Neal, 1985; Malchow et al., 1990; Akopian et al., 1991; DeVries and Schwartz, 1992; Jonz and Barnes, 2007; Sun et al., 2017; Country et al., 2019). Unless otherwise stated, all reagents and chemicals were sourced from Sigma-Aldrich (Oakville, ON, Canada). Eyes were removed and placed in cold Ca^2+^-free Ringer’s solution, containing (in mmol l^-1^) 120 NaCl, 2.6 KCl, 1 NaHCO_3_, 0.5 NaH_2_PO_4_, 1 sodium pyruvate, 4 Hepes and 16 glucose at pH 7.8. Whole retinas were placed in hyaluronidase (100 U ml^-1^, cat. no. H-3506) in L-15 solution for 20 min at room temperature. L-15 solution was comprised of 70% L-15 (Leibovitz’s medium) and 30% Ca^2+^-free Ringer’s. Retinas were washed 3 times for 3 min in L-15 solution and then placed in L-15 solution containing 7 U ml^-1^ papain (cat. no. 3126, Worthington Biochemical Corporation, Lakewood, WI, USA) for 40 min. Papain was activated by 2.5 mmol l^-1^ L-cysteine. Retinas were rinsed 3 times in L-15 solution. Small (∼4 mm^2^) sections of retina were removed and mechanically dissociated by gentle trituration in L-15 solution. The resulting cell suspension was plated and allowed to settle for up to 20 min.

### Morphometric analysis of neuronal subtypes

Phase contrast microscopy was performed on a Zeiss Axio Observer A1 microscope (Carl Zeiss, Oberkochen, Germany). Images were captured with QCapture (QImaging Corp., Surrey, BC, Canada) and measurements were performed with ImageJ (Schneider et al., 2012). All neurons were identified by characteristic stellate morphology, flat bodies, and thick dendrites that are characteristic of horizontal cells (Dowling et al., 1985; Tachibana, 1981). Morphometry of specific subtypes (i.e. H1, H2, H3 and H4) was performed following procedures described by Country et al. (2021), which identifies subtypes based on the ratio of dendritic field area to soma area (*r*_d/s_). Briefly, an ellipse representing the approximate dendritic spread was calculated using the formula for area (*A*):

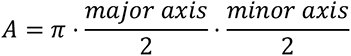

This process was repeated for the soma, and the *r*_d/s_ of the ellipse representing the dendritic field versus the ellipse representing the soma was measured in all cells. This analysis was performed for cells isolated in the summer and winter to determine whether any changes in size or shape could be observed.

### Mass spectrometry-based lipidomics

Lipidomics was performed at the Carleton Mass Spectrometry Centre, Carleton University, Ottawa, Canada. For sample preparation, whole retinas were removed from goldfish (as above) in July or January 2017, placed in PBS in centrifuge tubes and frozen immediately in liquid N_2_. Retinas were combined in a 15 ml centrifuge tube (Corning) and 1 ml of 2% acetic acid in methanol (AcMeOH) was added. Tissue was sonicated on ice using a 130 W ultrasonic processor for 3 min with the pulse on for 20 s and pulse off for 10 s using 50% amplitude.

Lipid extraction was accomplished using a method modified from Bligh and Dyer (1959). Briefly, each sonicated sample was transferred to a 10 ml glass Kimble tube containing 3.2 ml of 0.22 µm filtered 0.1 mol l^-1^ Na-acetate via triplicate rinses with 1 ml of 2% AcMeOH. 60 μl of a 1 mmol l^-1^ C13:0 lysophosphatidylcholine solution in ethanol (EtOH) was added as an internal standard. 3.8 ml of chloroform was then added and the tubes were vortexed for 30 s followed by 2 min centrifugation at 2,000 rpm and 4°C. The bottom phase was collected with a Pasteur pipette and transferred to a new 10 ml glass Kimble tube. Extractions on the aqueous phase were repeated 3 more times in the same manner using 2 ml of chloroform and combined. The combined chloroform extract was evaporated under a stream of N_2_ gas until a dark yellow oily residue remained. The residue was then resuspended in 600 μl of 10 mmol l^-1^ ammonium acetate in absolute EtOH. The samples were then transferred to an amber glass tube, flushed with N_2_ gas and stored in a –20°C freezer until analysis within a week. Samples requiring longer storage times were stored at –80°C.

Glycerophospholipid samples were separated via high performance liquid chromatography (HPLC) and analyzed by mass spectrometry (MS) and tandem mass spectrometry (MS^2^). C_4_ reversed phase microflow chromatography columns were constructed in-house as previously described (Canez et al., 2016). Lipids were separated using a ternary gradient program consisting of 10 mmol l^-1^ ammonium acetate in 30% methanol/70% MilliQ water (solvent A), 10 mmol l^-1^ ammonium acetate in 75% isopropanol (IPA)/25% methanol (solvent B), and methyl tert-butyl ether, MTBE (solvent C). The gradient began at 100% A, increased linearly to 100% B by 41.4 min, then to 100% C by 48 min before returning to 100% A between the 59 and 60 min time points. An average flow rate of 135 µl min^-1^ was used and each analytical run was followed by a full blank run to further clean the system and ensure no lipid carryover occurred.

For quantitative analysis, 12.5 μl lipid extract was diluted with 37.5 μl of 10 mmol l^-1^ ammonium acetate, separated using a Dionex Ultimate 3000 HPLC (Thermo Fisher Scientific, Waltham, MA, USA) in the manner described above and introduced into a QTRAP 4000 hybrid triple quadrupole linear ion trap mass spectrometer (AB Sciex, Framingham, MA, USA) using electrospray ionization. Triplicate analyses for both the winter and summer samples were performed in positive ion mode according to the parameters outlined in the Supplementary Information (Table S1).

**Table 1.** Species of phosphatidylethanolamine (PE) from retinal tissue detected using low resolution mass spectrometry that increased in winter. Possible structures of these lipids are derived from the lipid database on LIPID MAPS Lipidomics Gateway and are indicated based on the nominal mass of each lipid using IUPAC-IUBMB lipid nomenclature (total number of acyl chain carbons:total number of double bonds).

| Low resolution protonated masses of PE species (m/z) | Possible structures |
| --- | --- |
| 688 | PE(32:2), PE(O-33:2), PE(P-33:1) |
| 704 | PE(33:1), PE(O-34:1), PE(P-34:0) |
| 730 | PE(35:2), PE(O-36:2), PE(P-36:1) |
| 738 | PE(36:5) |
| 820 | PE(42:6) |

In order to increase the granularity of our dataset, vinyl-ether containing lipids (plasmalogens) were distinguished from alkyl-ether containing lipids via reaction with formic acid, as described by Fhaner et al. (2012). Formic acid-treated lipid samples were redissolved in 10 mmol l^-1^ ammonium acetate in EtOH for MS analysis, as described above. To reveal greater qualitative detail, individual lipids with statistically significant concentration differences between summer and winter were re-analyzed at high resolution using a 1290 HPLC coupled to a 6550 Q- TOF mass spectrometer (Agilent, Santa Clara, CA, USA). The HPLC solvents and conditions were the same as described above and the MS was run in positive ion mode for high mass accuracy determination, and in negative ion mode using MS^2^ with a collision energy of 35 eV for fatty acyl group identification.

For statistical analyses, Analyst 1.5.1 (AB Sciex) was used to generate extracted ion chromatograms, which were inputted into MultiQaunt 2.1.1 (AB Sciex) to calculate peak areas. To standardize data, the area sum for every extracted ion was calculated for winter and summer samples and normalized to each other. The individual peak areas were then standardized to the area of the internal standard. Technical replicates enabled the average area with standard deviation to be calculated for each ion. To identify significant alterations in abundance between the winter and summer samples, fold changes were calculated and a Student’s *t*-test was performed (α=0.05). To control the false discovery rate, the Benjamini-Hochberg method was applied with an α=0.1533 (Benjamini and Hochberg, 1995). P-values from the Student’s *t*-test and Benjamini-Hochberg method were compared. High resolution MS data were analyzed using MassHunter Qualitative Analysis B.07.00 (Agilent). MS files were converted to MGF format using Proteowizard’s MSConvert tool (Chambers et al., 2012) using the Threshold Peak Filter to select the 100 most intense peaks by counts; mzML format conversion without filters was also performed. The resulting MGF file was searched against the LipidBlast database (Kind et al., 2013) using NIST MSPepSearch software using the following settings: Library LipidBlast-neg, MinMF 100, HITS 3, MzLimits 1-2000, MinInt 1, OnlyFound, OutPrecursorMZ, OutDeltaPrecursorMZ, OutSpecNum, HiPri. Hits were then filtered by m/z to each of the 5 targeted lipid masses and compared to their extracted ion chromatograms using MZmine 2 (Pluskal et al., 2010). Annotating the targeted masses was performed by selecting database matches to PE species with retention times corresponding to the elution profile observed in the chromatograms of target masses. Annotations were confirmed using the NIST MS Search 2.0 program for head-to-tail comparisons to the LipidBlast database.

### Immunohistochemistry and confocal imaging

Live and fixed cell images were acquired with a Nikon (Tokyo, Japan) TiE A1-R high-speed resonant laser scanning confocal microscope using a 60*×*/NA1.2 objective lens. For cell fixation, 3.5% w/v paraformaldehyde was used in a 2% w/v sucrose in Dulbeco’s phosphate-buffered solution (DPBS, 311-425-CL, Wisent, St-Bruno, Québec, QC, Canada) by incubating cells for ∼15 min at room temperature, followed by a wash with DPBS 3 times. Cells were incubated with 0.5% v/v Triton X-100 in permeabilization buffer (20 mmol l^-1^ Hepes, 300 mmol l^-1^ sucrose, 50 mmol l^-1^ NaCl, 3 mmol l^-1^ MgCl_2_, diluted in distilled water) for 3 min. Cells were blocked with wash buffer (5% fetal bovine serum in DPBS) for a minimum of 1 h prior to incubation with antibodies. Monoclonal anti-α-tubulin produced in mouse (cat. no. T6074, Sigma-Aldrich) labelled microtubules and was used at a concentration of 1:200 and prepared in wash buffer. Following 3 wash steps with 5 min each in wash buffer, cells were stained with Alexa Fluor 488 rabbit anti-mouse IgG secondary antibody (cat. no. A11059, Invitrogen, Mississauga, ON, Canada) at a 1:200 dilution. Actin staining was simultaneously performed by incubation with Alexa Fluor 546 phalloidin (cat. no. A22283, Invitrogen) at 1:100 dilution.

DAPI (4′,6-diamidino-2-phenylindole, D1306, Invitrogen) staining was performed by incubation at 1:500 for 10 min in DPBS, following further rinse steps. All fixed cells were first seeded onto 35 mm dishes coated with Cell-Tak (Corning, Tewksbury, MA, USA). Stained images were captured as z-stacks and maximum intensity projections were generated using ImageJ (Schneider et al., 2012).

### Atomic force microscopy

Membrane stiffness (or elasticity) was measured in isolated neurons during the summer (June to September) and winter (January) months over the period of two consecutive years (2011*−*2013). Cell suspensions were allowed to adhere to 35 mm glass-bottom dishes (MatTek, Ashland, MA, USA) for at least 10 min. Live cells were first imaged by confocal microscopy, followed by nano-indentation using a NanoWizard II (JPK Instruments, Berlin, Germany) atomic force microscope (AFM). Cantilevers (MSCT-AUHW, Veeco, Plainview, NY, USA) were calibrated as previously described (Haase and Pelling, 2013). Nano-indentation was performed by recording a minimum of 10 force curves for each cell above the most central nuclear region (approach rate 2 Hz, 10 μm s^-1^, set-point of 1 nN). Force-distance curves were fit to the first 200 nm of deformation using the modified Hertz model for a conical tip using PUNIAS Software (Carl et al., 2001) to provide a measure of Young’s modulus (Pa). In some experiments, cells were pre-treated with Hoechst 33342 (Invitrogen) in order to label DNA so that nano-indentation measurements could be compared between measurements performed on- (N = 11) and off- nucleus (N = 11) for the same cell. Mean values for Young’s modulus were used for each cell (N > 10 force curves each) and overall statistics were calculated using a Student’s *t*-test with significance indicated by P < 0.05.

### Electrophysiology

Whole-cell, voltage-clamp recordings were obtained from neurons during the summer (June to September) and winter (January to March) months of 2005 and 2006 at Dalhousie University in Halifax, Canada. Electrodes were made from capillary glass (cat. no. 2502, Chase Scientific Glass, Inc., Rockwood, TN, USA) and pulled on a vertical puller (Model 730, David Kopf Instruments, Tujunga, CA, USA). Electrodes had a tip resistance of 5–8 M*Ω* when filled with intracellular recording solution containing (in mmol l^-1^): 120 KCl, 10 NaCl, 0.5 CaCl_2_, 2 Mg-ATP, 5 EGTA, 10 Hepes, pH adjusted to 7.4 with KOH. Extracellular recording solution contained (in mmol l^-1^): 120 NaCl, 5 KCl, 2.5 CaCl_2_, 2 MgCl_2_, 10 Hepes, 10 glucose (pH adjusted to 7.8 with NaOH). With these solutions the calculated equilibrium potential for K^+^ was –80.5 mV. Liquid junction potentials (*V*_L_) of 4.2 mV were calculated using pCLAMP software (Axon Instruments, Sunnyvale, CA, USA) and subtracted from pipette potentials (*V*_p_) to determine actual membrane potential (*V*_m_), according to the equation: *V*_m_ = *V*_p_ − *V*_L_. Current– voltage data were corrected for *V*_L_.

Voltage-clamp protocols were performed using an Axopatch-1D amplifier (Axon Instruments) and pCLAMP software. Recorded signals were converted using a DigiData 1200 interface (Axon Instruments). Cells were plated in 35 mm plastic Petri dishes (Nunclon, Nunc A/S, Roskilde, Denmark). From a holding potential of −60 mV, currents were evoked by changing *V*_m_ to a series of test potentials by a voltage-ramp protocol, in which the membrane potential was continuously changed (0.18–0.22 mV ms^−1^) over a period of 1 s. Signals were filtered at 2 kHz and digitized at a rate of 10 kHz. Membrane capacitance (*C*_m_) was measured using pCLAMP.

Reversal potential (*V*_rev_) was measured directly from *I-V* curves of winter neurons as an estimate of their resting membrane potential (see Discussion). In summer neurons, however, it was not possible to obtain accurate values of *V*_rev_ because of the flat slope conductance, i.e. high input resistance (*R*_in_), within the physiological range in these cells (see Fig. 5). To compare membrane activity of winter versus summer neurons, we therefore measured ion channel current in winter neurons at *V*_rev_ (which was zero) and compared this with current measured in summer at the same membrane potential. In some experiments, the inverse slope of the *I–V* relationship was used to calculate *R*_in_ from each cell by measuring the change in current from a stable region (of up to 40 mV) within the voltage range of *−*80 mV to *−*15 mV. *R*_in_ was then calculated using Ohm’s Law: *R*_in_ = *V* / *I*. All data were analysed using pCLAMP software and figures were arranged using Prism 5.0 (GraphPad Software Inc., La Jolla, CA, USA). Student’s *t*-tests were performed using Prism 5.0.

### Microspectrofluorometry (intracellular Ca^2+^ imaging)

Data were obtained from neurons during the summer (August and September) and winter (January and February) months between 2016*−*2020. Relative changes in free intracellular Ca^2+^ concentration ([Ca^2+^]_i_) were measured by microspectrofluorometric imaging following Country et al. (2019). Isolated cells were plated in 35 mm dishes (Corning) fitted with perfusion chambers (Warner Instruments Inc, Hamden, CT, USA, cat. no. RC-33DL). Dishes were pre-coated with 0.01% poly-L-lysine (cat. no. A-005-C). Cells were protected from light and incubated in extracellular solution containing (in mmol l^-1^) 120 NaCl, 5 KCl, 2.5 CaCl_2_, 2 MgCl_2_, 10 Hepes, and 10 glucose, with pH adjusted to 7.8 with NaOH. Added to this solution was 5 μmol l^-1^ membrane-permeant Fura-2 (Fura-2-LeakRes AM; Teflabs, Austin, TX, USA) and 0.1% (vol/vol) of a 10% (wt/vol) Pluronic F-127 solution for 30 min at room temperature.

Cells were washed three times to remove remaining esterified products. The chamber was continuously perfused at ∼1 ml min^-1^ by gravity-fed recording solutions at room temperature. Recording solution was removed from the chamber at the same rate using a variable-flow pump (Thermo Fisher Scientific).

Fluorescence imaging was performed with a Lambda DG-5 wavelength changer (Sutter Instruments, Novato, CA, USA) and a Chroma 79001 filter set (340 nm and 380 nm band-pass filters for excitation, 510 nm bandpass for emission; Chroma Technology, Bellows Falls, VT, USA). Excitation and emission light was passed through a 40*×* water-immersion objective lens (MRF07420 CFI Fluor, Nikon) optimized for UV transmission on an upright microscope (FN-1, Nikon). Images were collected with a CCD camera (QImaging) by focusing on a region of interest surrounding a soma. Excitation wavelength was changed between 340 nm and 380 nm and emission intensity was recorded for both excitation wavelengths every 2 s with NIS Elements software (Nikon). Imaging data were logged in Excel (Microsoft Corp., Redmond, WA, USA).

Raw traces from Ca^2+^ imaging experiments are presented as the ratio of fluorescence emission intensity after excitation with 340 nm and 380 nm (F_340_/F_380_). These values are proportional to [Ca^2+^]_i_. Parameters chosen to describe each spontaneous action potential event included event frequency, duration, amplitude, area under the curve (AUC), and time to peak (TTP) from baseline level. Events were analysed using a peak analysis algorithm in OriginPro 2016 (OriginLab Corp. Northampton, MA, USA). Frequency was measured as the number of events per sampled time period (5 min). Duration was measured as the time (s) between the start and end of each event using a local maximum function compared to baseline. Amplitude (F_340_/F_380_) was measured as the difference between the peak value and baseline directly before the event, as determined by the Origin peak analysis gadget using a second derivative algorithm for determining baseline values. AUC (F_340_/F_380_ · s) approximates Ca^2+^ influx into the cytosol and was calculated as the exact integral of amplitude over time of each event from [Ca^2+^]_i_ baseline:

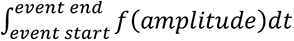

TTP was the time (s) between the start of the event and the peak amplitude of that event. For each cell that presented action potentials during a 5 min sampling period, duration, amplitude, AUC, and TTP are reported as the average of all events within that period. Action potentials were only included if their peaks fell within the 5 min sampling period, and if their durations were ≤ 300 s. Data were analyzed with the Student’s *t*-test using Prism 7 (GraphPad Software Inc.).

## RESULTS

### Cell morphology and membrane surface area were consistent between seasons

Neurons (i.e. horizontal cells) isolated from goldfish retina were routinely identified by their large, stellate morphology and thick dendrites. We identified all subtypes of horizontal cells (Fig. 1A–D) based on morphological description from previous studies (Dowling et al., 1985; Tachibana, 1981; Country et al., 2021) and observed no apparent differences in morphology between summer and winter. Plotting the distribution of the ratio of dendritic field area to soma area provided an estimate of the subtypes present in the dissociation (Country et al., 2021). Calculation and plotting of the *r*_d/s_ values of 80 neurons showed that the distribution of cells isolated during the summer and winter were similar, with median *r*_d/s_ values of 7.2 and 6.2, respectively (Fig. 1E). Analysis with the Mann-Whitney *U* test indicated that *r*_d/s_ distributions were not significantly different from each other (P = 0.20). Moreover, during both seasons the dominant subtypes observed were H1 and H2 with *r*_d/s_ values of less than 11, while H3 and H4 subtypes, with higher *r*_d/s_ values, were fewer in number (Fig. 1E). Mean membrane capacitance (*C*_m_) of neurons was measured from electrophysiological experiments to estimate cell size (Fig. 1F). Biological membranes have a specific capacitance of 0.01 pF µm^-2^ (Hille, 2001). *C*_m_ was therefore used to estimate total membrane surface area. *C*_m_ was 26.85 ± 1.6 pF in summer (N = 46) and 26.90 ± 1.1 pF in winter (N = 81), indicating that neuronal membrane surface area did not differ between seasons (P = 0.49, Student’s *t*-test).

**Fig. 1.**
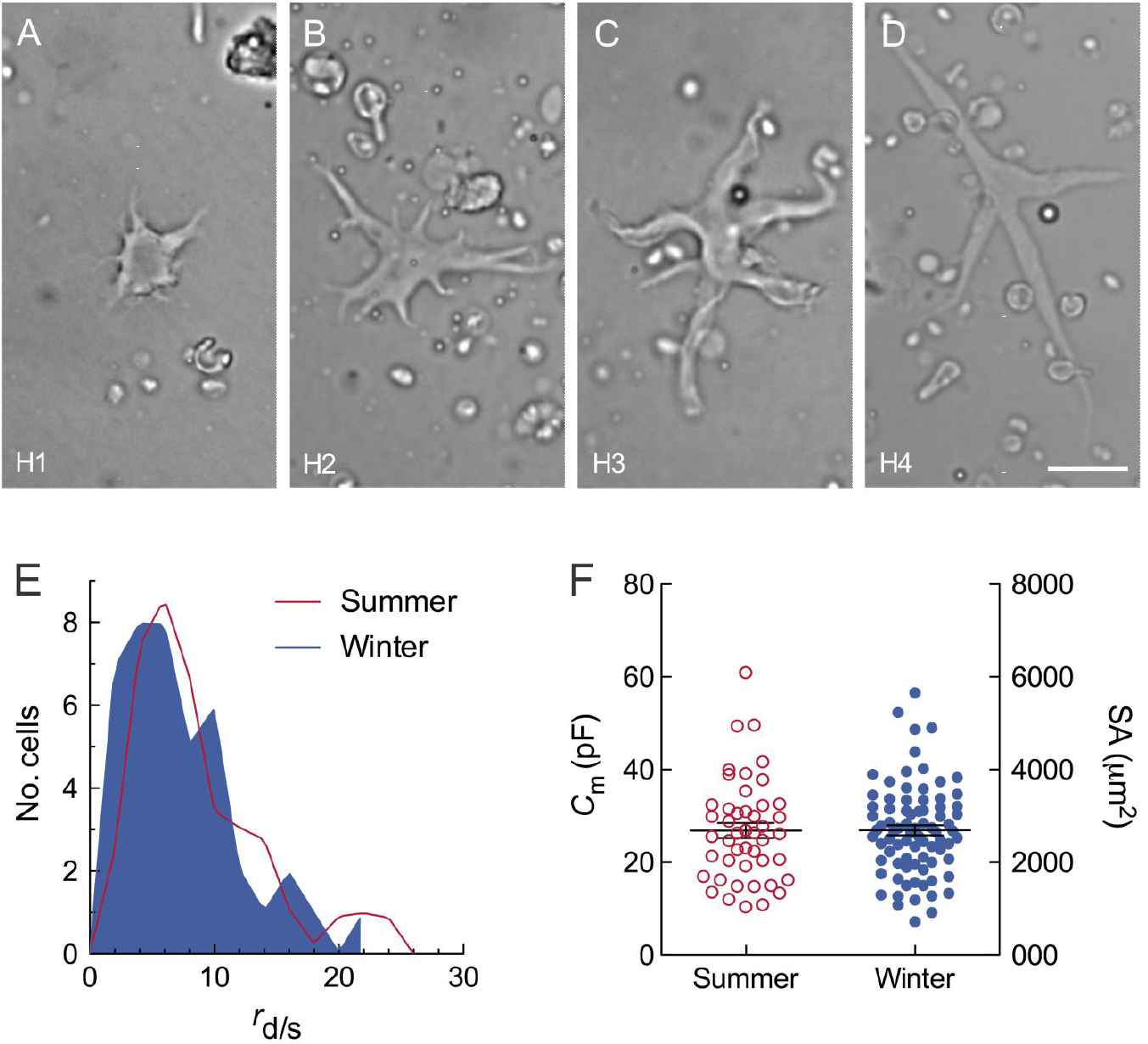
Morphology and subtype-specific distribution of neurons (horizontal cells) displayed no observable differences between seasons. (A–D) Examples of subtypes, H1, H2, H3 and H4, isolated from goldfish retina and identified using phase-contrast microscopy. Scale bar in D = 10 µm and applies to all panels. (E) Frequency distribution of the ratio of dendritic field area to soma area (*r*_d/s_) estimated the prevalence of subtypes isolated during the summer (line) and winter (shaded area). Summer, median *r*_d/s_ of 7.2 and interquartile range of 5.0–11.6; Winter, median *r*_d/s_ of 6.2 and interquartile range of 3.7–9.6. Lines were fit by local regression using locally weighted scatterplot smoothing. (F) Mean ± s.e.m. membrane capacitance (*C*_m_) in neurons isolated in the summer (open circles) and winter (closed circles), as calculated from electrophysiological experiments. *C*_m_ (left axis) is proportional to membrane surface area at an approximate rate of 0.01 pF µm^-2^ and was used to estimate cell size. Calculated surface area (SA) is shown in µm^2^ on the right axis.

### Winter neurons had a higher concentration of unsaturated phosphatidylethanolamines

Analysis of low resolution MS data yielded a total of 153 lipid species in goldfish retina that were above the limit of quantification, calculated using established protocols (Canez et al., 2016). Of these, 90 were identified as phosphatidylcholine (PC), 37 as PE, 19 as sphingomyelin (SM) and 7 as phosphatidylserine (PS) (Fig. 2A). Measurement of standardized area values from MS data and analysis using the Student’s *t*-test with correction for false discovery rate using the Benjamini-Hochberg method revealed that 5 PE species had significantly higher concentrations in winter (P < 0.05; Fig. 2B). These PE species are listed in Table 1 along with their proposed identities based on their nominal masses using IUPAC-IUBMB lipid nomenclature (total number of acyl chain carbons:total number of double bonds). PEs were unsaturated and detected at concentrations 1.8 to 2.7 times higher in the winter compared to summer, as illustrated in Figure 2B.

**Fig. 2.**
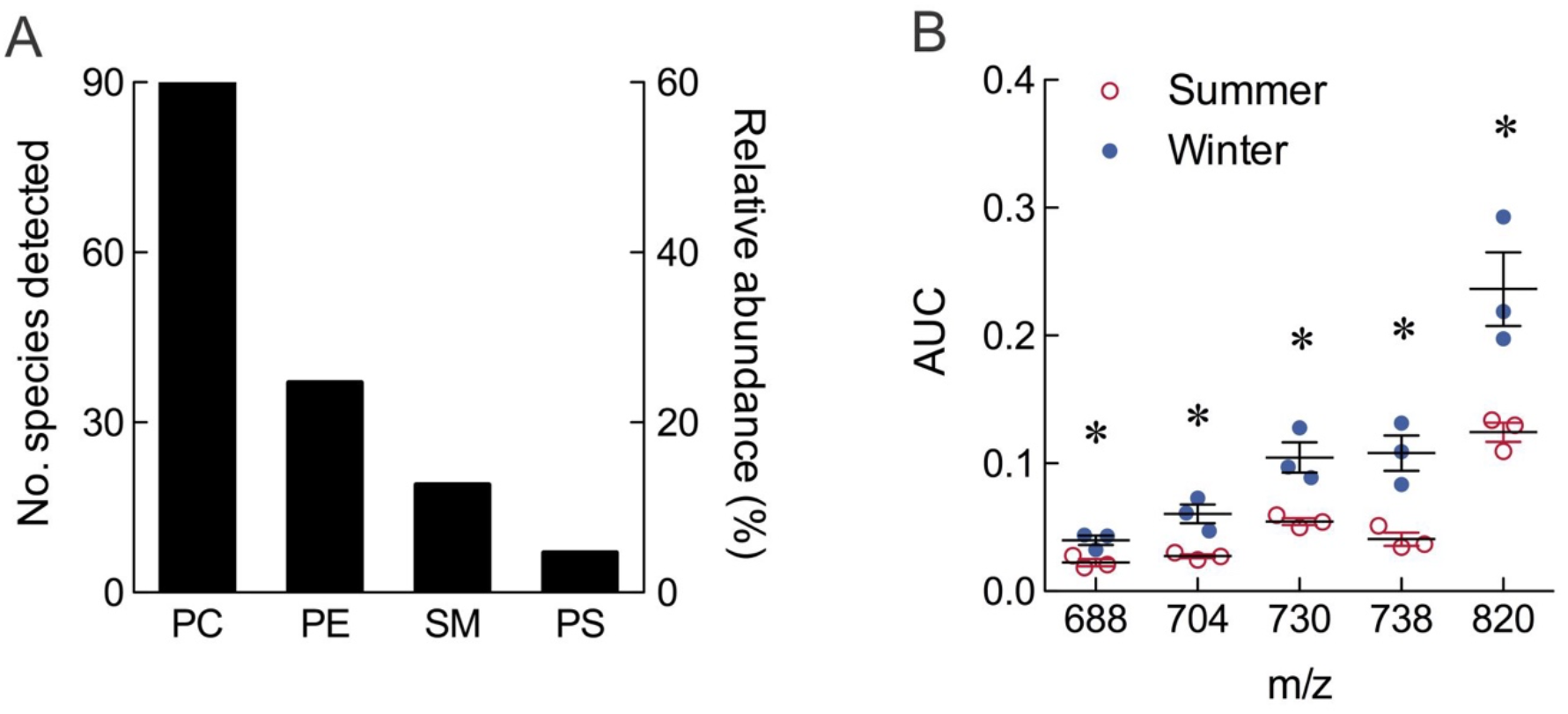
Mass spectrometry-based lipidomics of goldfish retina. (A) Number of species of phospholipids and their relative abundance, as determined by mass spectrometry. PC, phosphatidylcholine; PE, phosphatidylethanolamine; SM, sphingomyelin; PS, phosphatidylserine. (B) Changes in concentration of 5 species of PE between summer (open circles) and winter (closed circles). Area under the curve (AUC) was measured from mass spectrometry data and is shown for each mass-charge ratio (m/z). Asterisks indicate a significant difference between summer and winter using a Student’s *t*-test (*P* < 0.05) with correction for false discovery rate and the Benjamini-Hochberg method.

In order to further characterize the lipids that increased in abundance during winter, high resolution MS was conducted on these 5 PE species in positive ion mode to obtain accurate mass, and in negative ion mode to elucidate the specific fatty acyl composition of each via MS^2^. It was immediately discovered, based on accurate mass measurements, that one of the PE masses contained co-eluting diacyl- and ether-linked species that were indistinguishable in the quantitative results. In order to further characterize the ether-linked lipid, formic acid hydrolysis was performed (Fhaner et al., 2012) to determine whether it was a plasmanyl (vinyl-ether containing) species or plasmenyl (alkyl-ether containing) species. It was noted that both the plasmanyl and plasmenyl species were present at that mass and elution time at a ratio of 3:1, respectively. Table 2 shows the accurate masses of the PE species that increased in the winter months and reveals the relative percentages of the plasmanyl and plasmenyl lipids.

**Table 2.**
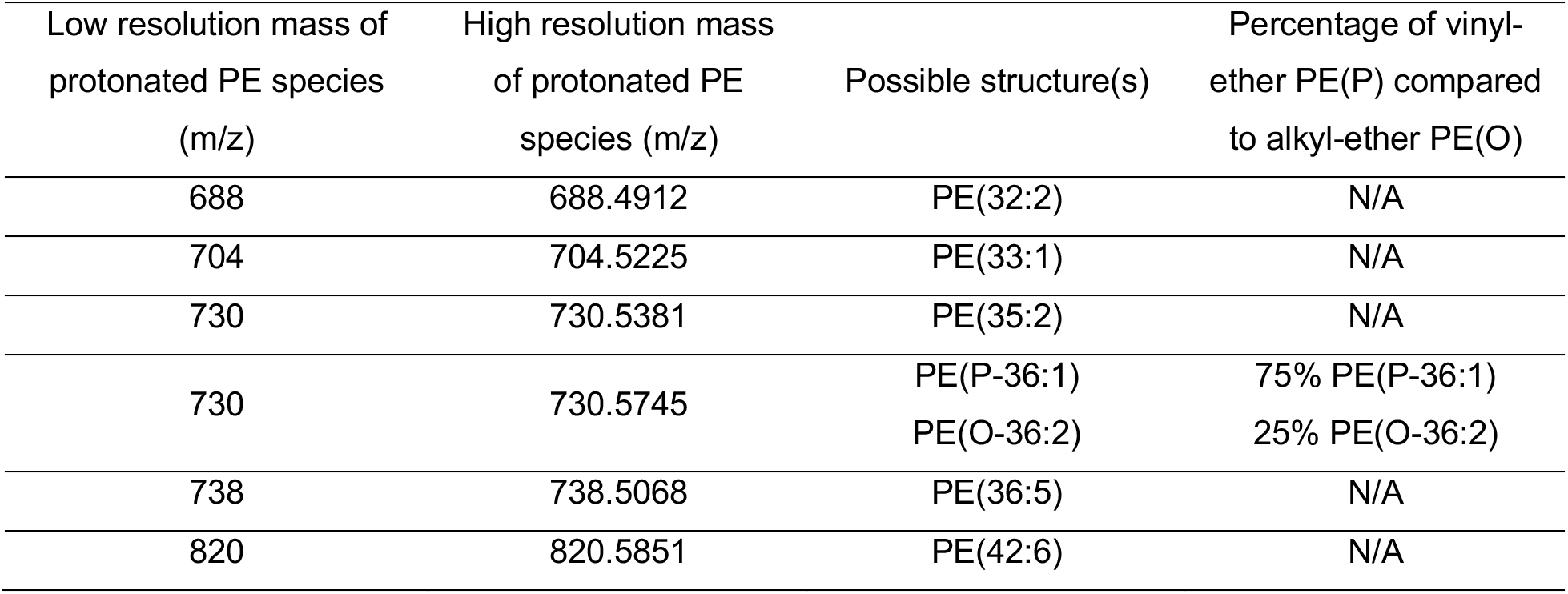
Accurate masses and identities of phosphatidylethanolamine (PE) that increased during winter. Species were obtained using high-resolution mass spectrometry, including the ratio of vinyl-ether to alkyl-ether species determined via formic acid hydrolysis.

| Low resolution mass of protonated PE species (m/z) | High resolution mass of protonated PE species (m/z) | Possible structure(s) | Percentage of vinyl-ether PE(P) compared to alkyl-ether PE(O) |
| --- | --- | --- | --- |
| 688 | 688.4912 | PE(32:2) | N/A |
| 704 | 704.5225 | PE(33:1) | N/A |
| 730 | 730.5381 | PE(35:2) | N/A |
| 730 | 730.5745 | PE(P-36:1)<br>PE(O-36:2) | 75% PE(P-36:1)<br>25% PE(O-36:2) |
| 738 | 738.5068 | PE(36:5) | N/A |
| 820 | 820.5851 | PE(42:6) | N/A |

Negative ion MS^2^ was conducted on all PE masses in Table 2 in order to obtain their identities. Lipid MS^2^ spectra were searched against the LipidBlast database using NIST MSPepSearch software and annotations were confirmed using the NIST MS Search 2.0 program via head-to-tail comparisons to the LipidBlast database. In negative ion mode, fatty acids are lost from lipid ions as both charged and neutral fragments, revealing the *sn* structural chemistry. The fatty acyl composition was deduced for the PE species by computationally analyzing the MS^2^ spectra and all PE species were identified in this manner with the exception of m/z 702.5069 due to interfering peaks in its MS^2^ spectrum. Lipids were identified based on the top scoring hit that was obtained and verified through manual inspection, as illustrated and described in Figure 3. Figure 3 includes the assumed positions of the double bonds and *sn* regiochemistry based on lipids reported in the NIST database. Unfortunately, determining *sn* regiochemistry or the position of the double bonds on the alkyl chains is beyond the capabilities of our analytical methodologies and the degree to which we are able to confidently identify the PE species is accurately reported in Table 3.

**Fig. 3.**
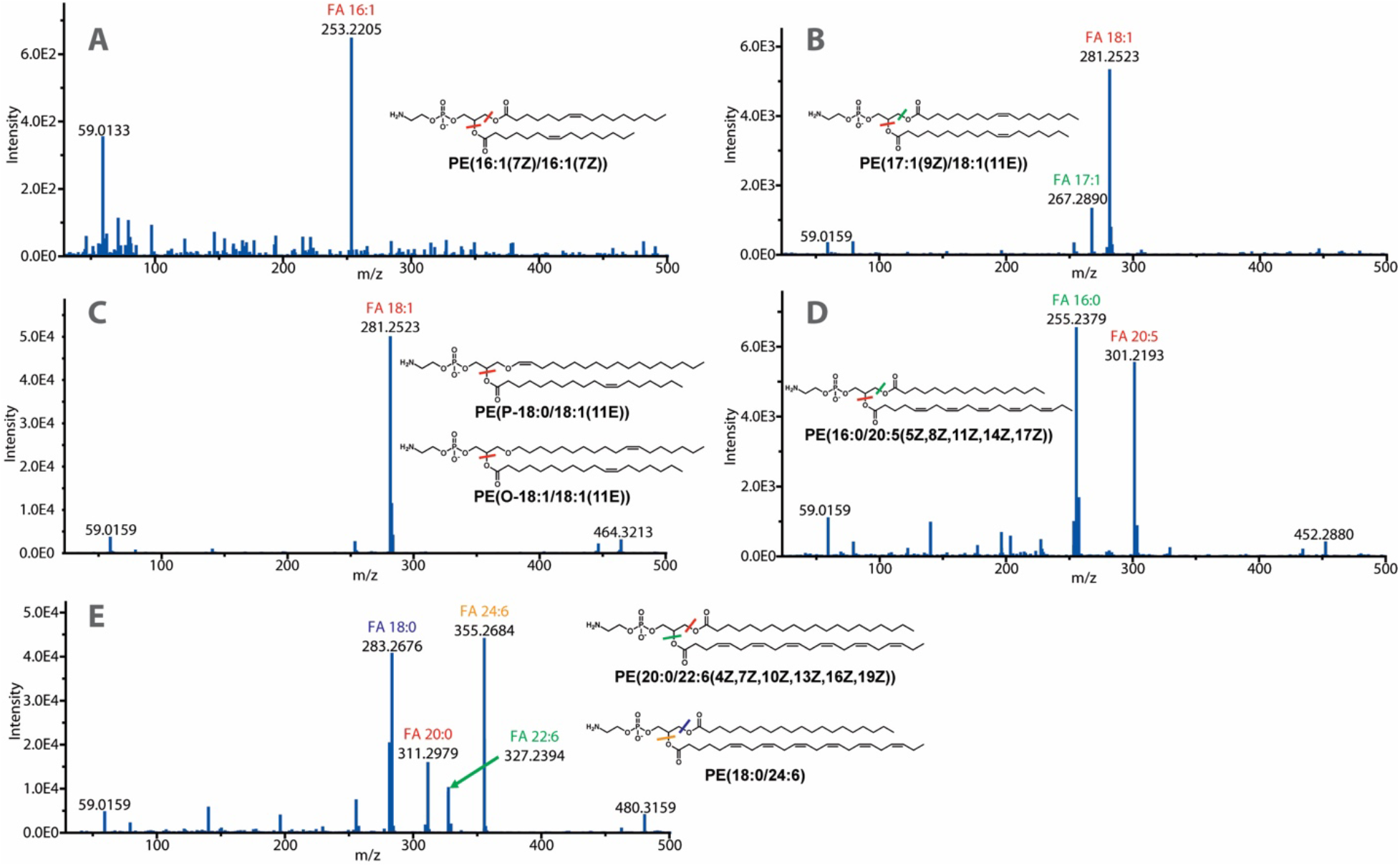
Tandem mass spectrometry analyses of statistically relevant PE species. (A) Negative ion MS^2^ of PE species at m/z 686.4845 identifies PE(16:1_16:1) via a single fatty acid fragment ion corresponding to 16:1. Red lines indicate sites of fragmentation. (B) Negative ion MS^2^ of PE species at m/z 728.5294 identifies PE(17:1_18:1) via two fatty acid fragments. Fragmentation sites are indicated by red and green lines for the identification of fatty acids 18:1 and 17:1, respectively. (C) Negative ion MS^2^ of PE species at m/z 728.5668 confirms the identification of PE(P-18:0_18:1) as well as PE(O-18:1_18:1) through the observation of an 18:1 fatty acid fragment ion. The red line indicates the site of fragmentation. Fragmentation will only occur at alkyl groups, therefore both PE(P) and PE(O) are expected to form one peak at m/z 281.2523 corresponding to the fatty acid 18:1. (D) Negative ion MS^2^ of PE species at m/z 736.5010 confirms the identification of PE(16:0_20:5) via two fatty acid fragments. Fragmentation sites are indicated by red and green lines for the identification of fatty acids 16:0 and 20:5, respectively. (E) Negative ion MS^2^ of PE species at m/z 818.5821 reveals the identities of two different co-eluting isobaric lipids, PE(20:0_22:6) and PE(18:0_24:6) through the observation of four different fatty acid fragment ions. Fragmentation sites are indicated by coloured lines which correspond to the colour of the fatty acid peak labels. The PE(20:0_22:6) species was determined to be approximately 25% as abundant as the PE(18:0_24:6) species based on fragment ion peak intensities. The PE structures in panels A through E are illustrated in an unambiguous manner in terms of double bond positioning and *sn* regiochemistry and are based on previously reported structures in the LipidBlast database. Methodological limitations prevent confirmation of this structural information and therefore the illustrated structures represent the most likely candidates based on known PE species in the literature.

**Table 3.** Identities of phosphatidylethanolamine (PE) species that increased in winter based on negative ion tandem mass spectrometry (MS^2^) analyses.

| Low resolution<br>mass of<br>protonated PE<br>species (m/z) | High resolution<br>mass of<br>deprotonated<br>PE species<br>(m/z) | Nominal structure | Full identification |
| --- | --- | --- | --- |
| 688 | 686.4845 | PE(32:2) | PE(16:1_16:1) |
| 704 | 702.5069 | PE(33:1) | Inconclusive - interference in MS<sup>2</sup> spectrum |
| 730 | 728.5294 | PE(35:2) | PE(17:1_18:1) |
| 730 | 728.5668 | 75% PE(P-36:1)*<br>25% PE(O-36:2)* | PE(P-18:0_18:1)<br>PE(O-18:1_18:1) |
| 738 | 736.5010 | PE(36:5) | PE(16:0_20:5) |
| 820 | 818.5821 | PE(42:6) | 75% PE(18:0_24:6)**<br>25% PE(20:0_22:6) |

\*Percentages determined through formic acid hydrolysis.
\*\*Structure not in database; determination through manual interpretation.

### Atomic force microscopy revealed an increase in membrane stiffness during winter

We used AFM to investigate whether membrane stiffness (or elasticity) differed between cells isolated during the summer versus winter months. Labelling of the cytoskeleton and nucleus was first performed to assess the morphology and relative location of these structures in neurons for AFM experiments. Microtubules and actin filaments were labelled with anti-α- tubulin and phalloidin, respectively, whereas the nucleus was labelled with DAPI (Fig. 4A).

**Fig. 4.**
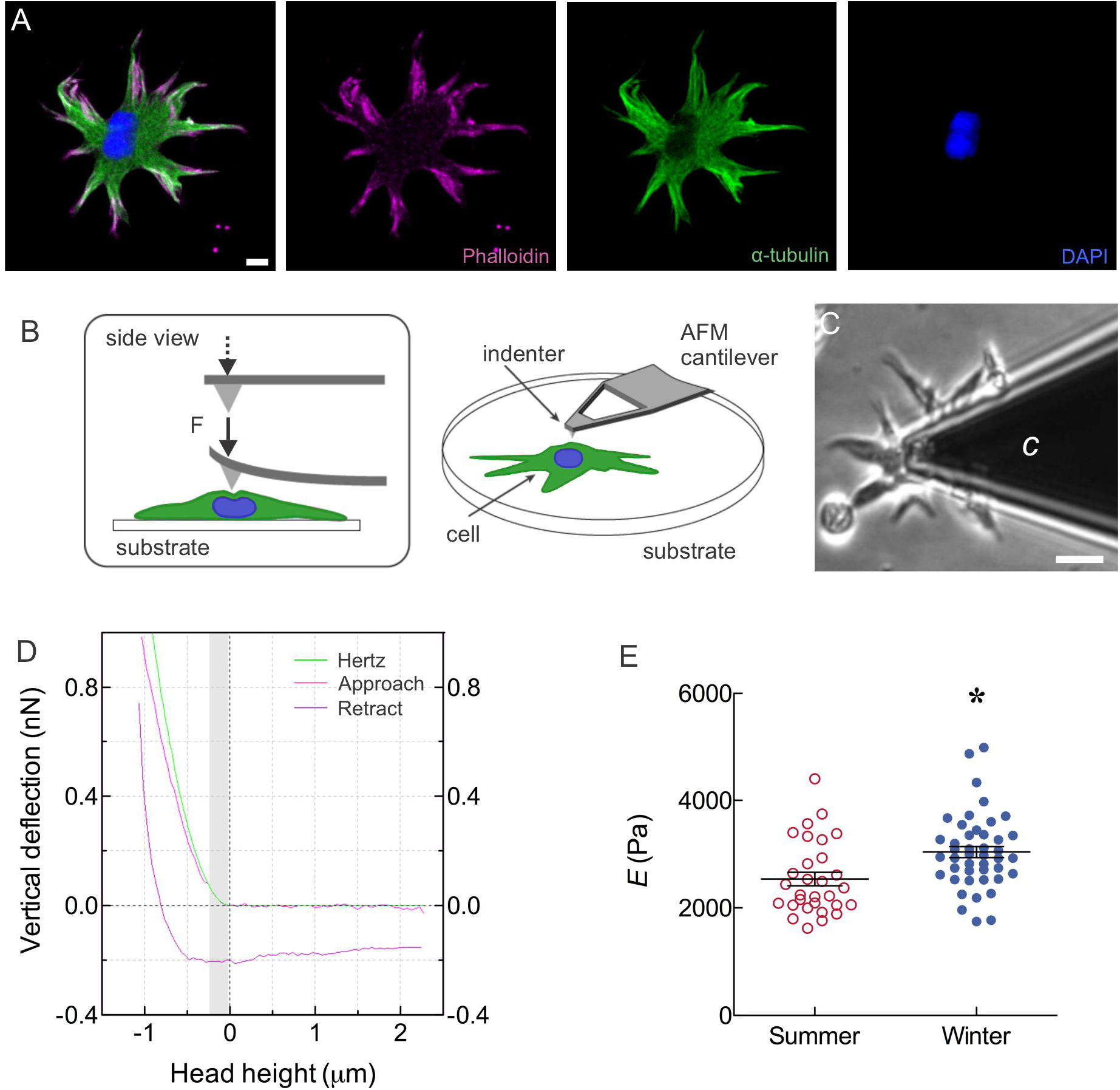
Membrane deformation by atomic force microscopy (AFM) demonstrated a seasonal change in membrane stiffness. (A) Neurons (horizontal cells) were identified by a large soma with a higher concentration of tubulin, and an extensive dendritic field containing a higher concentration of actin. The cell illustrated was labelled with phalloidin (magenta) to indicate actin filaments, anti-*α*-tubulin (green) to show microtubules, and DAPI (blue) to identify nuclei. All markers are shown in the panel at left, and individual markers are shown in subsequent panels. Scale bar = 5 µm. (B) Cartoon depiction of force nano-indentation measurements performed using an AFM cantilever and a low set-point force (F) of 1 nN applied above the central region of a cell on a glass substrate. Modified from Haase and Pelling (2015). (C) Compression of the membrane of a neuron at the soma by a cantilever (*c*) (see Movie 1 in the Supplementary Information). Scale bar = 10 µm. (D) An exemplar force curve demonstrates the fit (green) to the Hertz model of 200 nm from the point of contact (indicated by grey vertical bar). Approach (magenta) and retract (violet) curves are also shown. Image generated from one indentation curve using JPKSPM data processing software (see Materials and Methods). Curves are shown with smoothing. (E) Mean ± s.e.m. Young’s modulus (*E*) showing an increase in membrane stiffness in cells isolated during the winter (closed circles) compared to summer (open circles). Asterisk indicates a significant difference between groups (Student’s *t*-test, P = 0.003; N in summer = 30, N in winter = 44).

Using confocal microscopy, we found that neurons were often binucleated, with nuclei off-centre within the soma. Neurons possessed an extensive cytoskeletal system with a higher concentration of α-tubulin localized to the soma, and phalloidin labelling localized primarily to the dendrites.

AFM was employed to measure changes in apparent stiffness of neurons across sequential summer and winter seasons. Isolated neurons were allowed to settle onto glass-bottomed dishes, then imaged using a laser-scanning confocal microscope prior to nano- indentation via AFM. Force-indentation measurements were performed by positioning the probe over the largest central region of neurons (Fig. 4B). By initiating a low set-point value of 1 nN (maximum force applied), a conical tipped AFM cantilever was used to deform cell membranes at a frequency of 2 Hz (Fig. 4B,C; see Movie 1 in the Supplementary Information). Force-indentation curves were acquired and fit (the first 200 nm indentation) to a modified Hertz contact model in order to interpret the Young’s modulus (apparent stiffness) of the cells (Fig. 4D). Mean ± s.e.m. Young’s modulus for neurons dissociated in summer (2,537 ± 125 Pa, N = 30) was significantly less than in winter (3,041 ± 104 Pa, N = 44), as determined by the Student’s *t*-test (P = 0.003; Fig. 4E). An additional experiment employed Hoechst 33342 (a live-cell nuclear dye) to confirm that there was no difference in stiffness between measurements made above or adjacent to nuclei. Measurements were performed on- and off-nucleus on the same cells and resulted in no difference in measured stiffness (P = 0.36, Student’s paired *t*-test, N = 11).

### Neurons had reduced membrane excitability in winter

Voltage-clamp recordings were obtained from 23 cells isolated during the winter and 33 cells isolated during the summer. An *I–V* curve (Fig. 5A) was generated from these currents and displayed a similar profile as described in previous studies (Tachibana, 1983; Shingai & Christensen, 1986; Jonz and Barnes, 2007). Currents recorded from neurons were characterized by prominent inward rectification at negative potentials carried by inwardly-rectifying K^+^ channels, a sustained inward Ca^2+^ current through L-type channels that was activated near −40 mV, and outward rectification at potentials above +40 mV (Fig. 5A). These currents and ion channels have been described elsewhere (Tachibana, 1983; Shingai & Christensen, 1986; Jonz and Barnes, 2007).

**Fig. 5.**
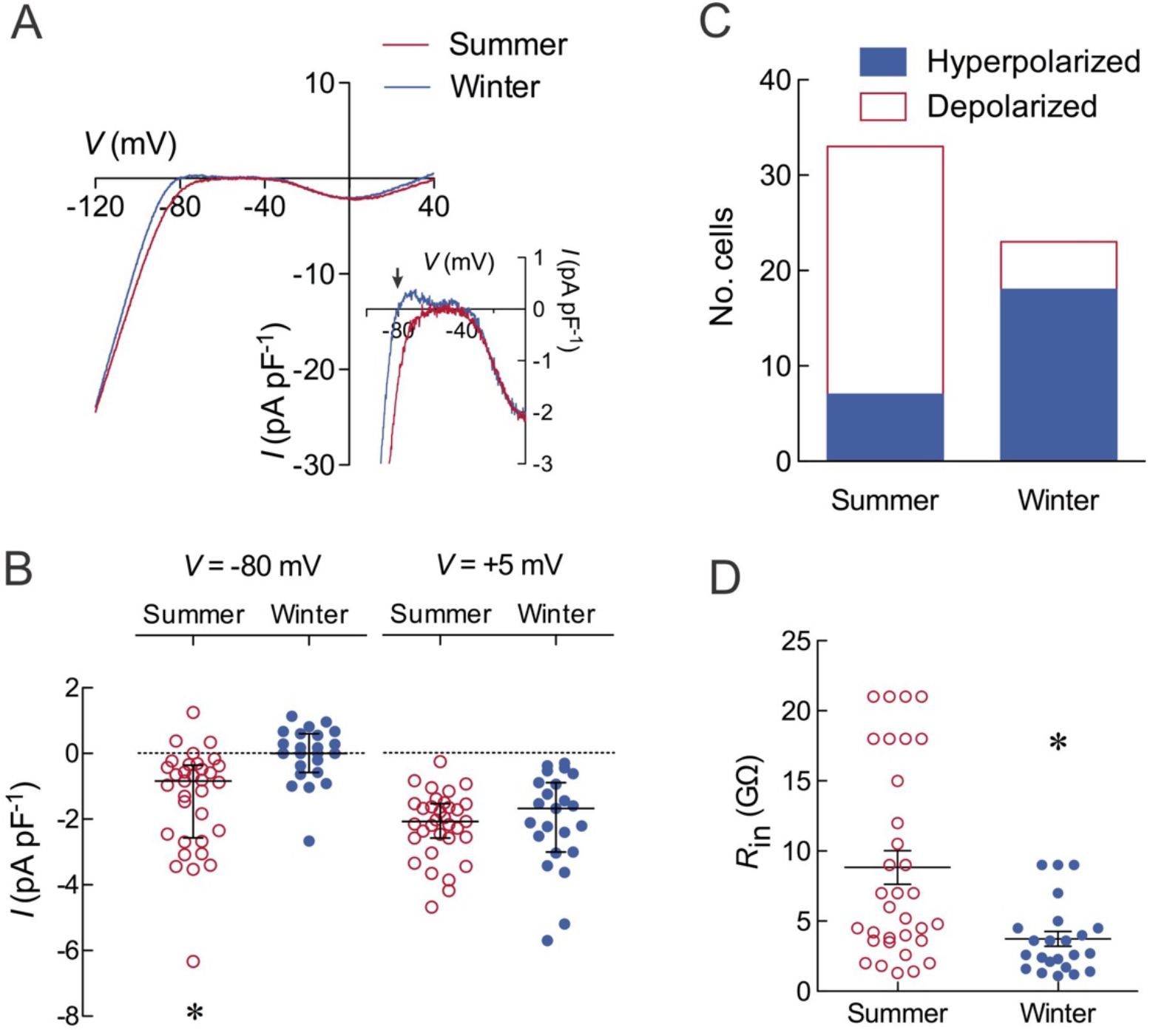
Neurons isolated during the winter had reduced membrane excitability. (A) Current-voltage (*I-V*) relationship from voltage-clamp electrophysiological recording. Membrane voltage of cells isolated during the summer (N = 33) or winter (N = 23) was changed progressively from –120 mV to +40 mV. Current density (pA pF^-1^) for each cell was calculated using membrane capacitance (*C*_m_) and the mean of all recordings is plotted. Inset, current density over the physiological range of potentials is shown at higher resolution. Arrow indicates the reversal potential (*V*_rev_ = *−*80 mV) in cells isolated during the winter. (B) Mean ± s.e.m. current density (pA pF^-1^) measured at *−*80 mV and +5 mV during voltage-clamp experiments. Asterisk indicates a significant difference between summer (open circles) and winter (closed circles) neurons (*P* = 0.0004, Student’s *t*-test). (C) Phenotype analysis of *I-V* curves demonstrated that cells isolated during the winter (shaded bars) were significantly (P < 0.0001) associated with a negative slope conductance and hyperpolarized *V*_rev_, compared to summer (open bars). (D) The input resistance (*R*_in_) for each cell in A was calculated following Ohm’s Law (*R*_in_ = *V* / *I*) for summer (open circles) and winter (closed circles). Asterisk indicates a significant decrease in *R*_in_ in cells isolated during the winter (*P* = 0.0007).

The reversal potential (*V*_rev_), where no net current was recorded, in winter neurons was measured as an averaged value from all traces and was –79.8 mV (Fig. 5A inset, arrow). This value was near the calculated equilibrium potential for K^+^, which was –80.5 mV. In summer neurons, *V*_rev_ was approximately –50 mV but could not be accurately determined because of the relatively flat slope conductance across the voltage axis. We therefore measured current density at –80 mV (to approximate *V*_rev_ in winter neurons) to compare differences in membrane activity between seasons. Compared to current measured in winter neurons, mean current density at –80 mV in cells isolated during the summer was significantly inward at –1.4 ± 0.3 pA pF^-1^ (Fig. 5B; Student’s *t*-test, P = 0.0004). This change in current density corresponded to the positive (depolarizing) shift in *V*_rev_ in cells isolated during the summer months (Fig. 5A). Current densities otherwise remained relatively unchanged between seasons at more negative potentials, and at potentials at which voltage-gated Ca^2+^ channels were active. At +5 mV, the approximate voltage at which peak Ca^2+^ current was recorded, there was no significant difference in current density between winter and summer neurons (Fig. 5B; Student’s *t*-test, P = 0.67).

From voltage-clamp data summarized in Figures 5A and B, we identified two different *I- V* phenotypes: (1) those from cells in which *I-V* curves displayed a negative slope conductance within the physiological range and an often hyperpolarized *V*_rev_, and (2) those from cells that did not display a negative slope conductance, had a relatively depolarized *V*_rev_, and inward current at –80 mV. Grouping cells according to these phenotypes, and analysis using the Fisher’s Test, demonstrated a significant association (P < 0.0001) between phenotype and season, such that cells with a negative slope conductance and hyperpolarized *V*_rev_ were more often observed in the winter (72%) compared to summer (28%), when the other phenotype was more common (Fig. 5C). In the retina, horizontal cells maintain a membrane potential in darkness of –35 mV to –20 mV and hyperpolarize when photoreceptors are exposed to flashes of light (Shingai and Christensen, 1986; Yang et al., 1988; Thoreson and Mangel, 2012; Sun et al., 2017). We therefore measured *R*_in_ from currents evoked between membrane potentials that correspond to resting levels in the intact retina. Cells isolated during the winter had a significantly lower mean *R*_in_ of 3.7 ± 0.5 G*Ω* compared to 8.8 ± 1.2 G*Ω* in the summer (Student’s *t*-test, P = 0.0007; Fig. 5D).

### Spontaneous Ca^2+^-based action potential activity decreased in winter

Based upon the changes in membrane excitability observed in neurons between summer and winter, we hypothesized that action potentials generated by these cells would be subject to seasonal variation. Spontaneous changes in [Ca^2+^]_i_ were measured in a total of 144 cells. Cells isolated in the summer typically displayed a plateau phase of the Ca^2+^-based action potential shortly after reaching peak [Ca^2+^]_i_, whereas cells isolated during the winter months displayed a profile characterized by a transient rise in [Ca^2+^]_i_ (Fig. 6).

**Fig. 6.**
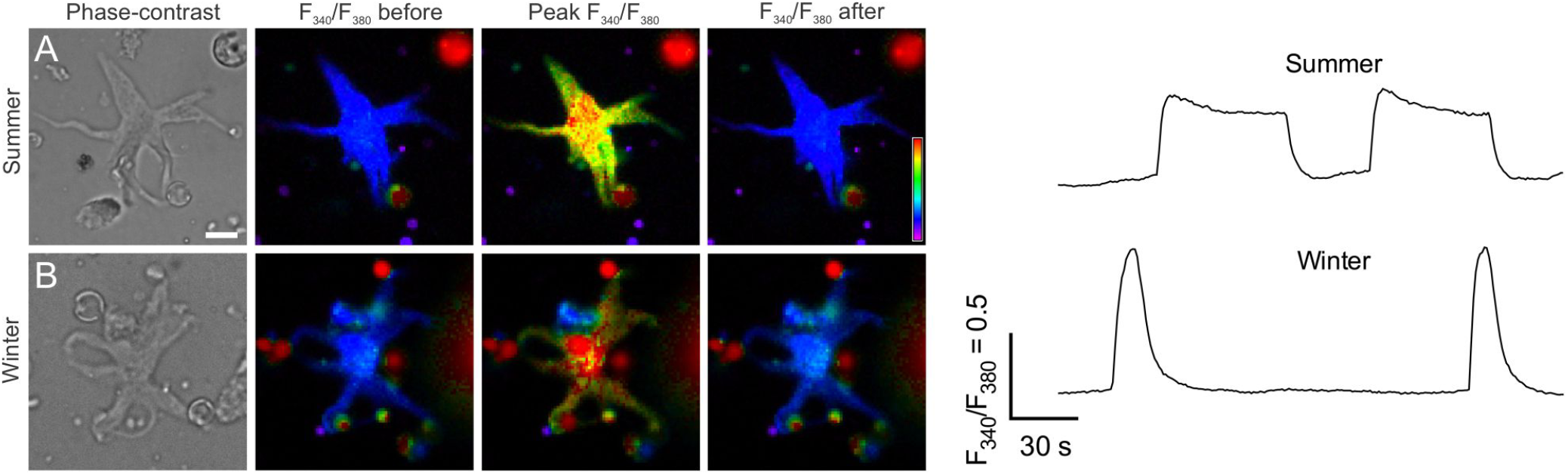
Microspectrofluorometry of spontaneous Ca^2+^-based action potentials in cells isolated during the summer and winter. Fluorescence imaging of changes in intracellular Ca^2+^ concentration ([Ca^2+^]_i_) in neurons using Fura-2 during summer (A) or winter (B). The fluorescence ratio (F_340_/F_380_) is shown before, during, and after the peak of action potentials. Fluorescence images correspond to the phase-contrast images of each cell at left. Scale bar = 10 µm and applies to all panels. A scale of relative change in fluorescence is shown. (Right) Changes in [Ca^2+^]_i_ with time in cells isolated during the summer (upper trace) and winter (lower trace). Action potentials during the summer were of longer duration. Scale bars indicate F_340_/F_380_, which is proportional to [Ca^2+^]_i_, and time in seconds.

To quantify seasonal changes in spontaneous [Ca^2+^]_i_ events, data from event amplitude, duration, AUC, frequency and time-to-peak were compiled from up to 208 spontaneous events from 76 cells in summer, and 209 spontaneous events from 68 cells in winter. Means ± s.e.m. for summary data are shown in Figure 7. Amplitude and AUC values were obtained from relative changes in the Fura-2 emission ratio (F_340_/F_380_) and are proportional to [Ca^2+^]_i_.

**Fig. 7.**
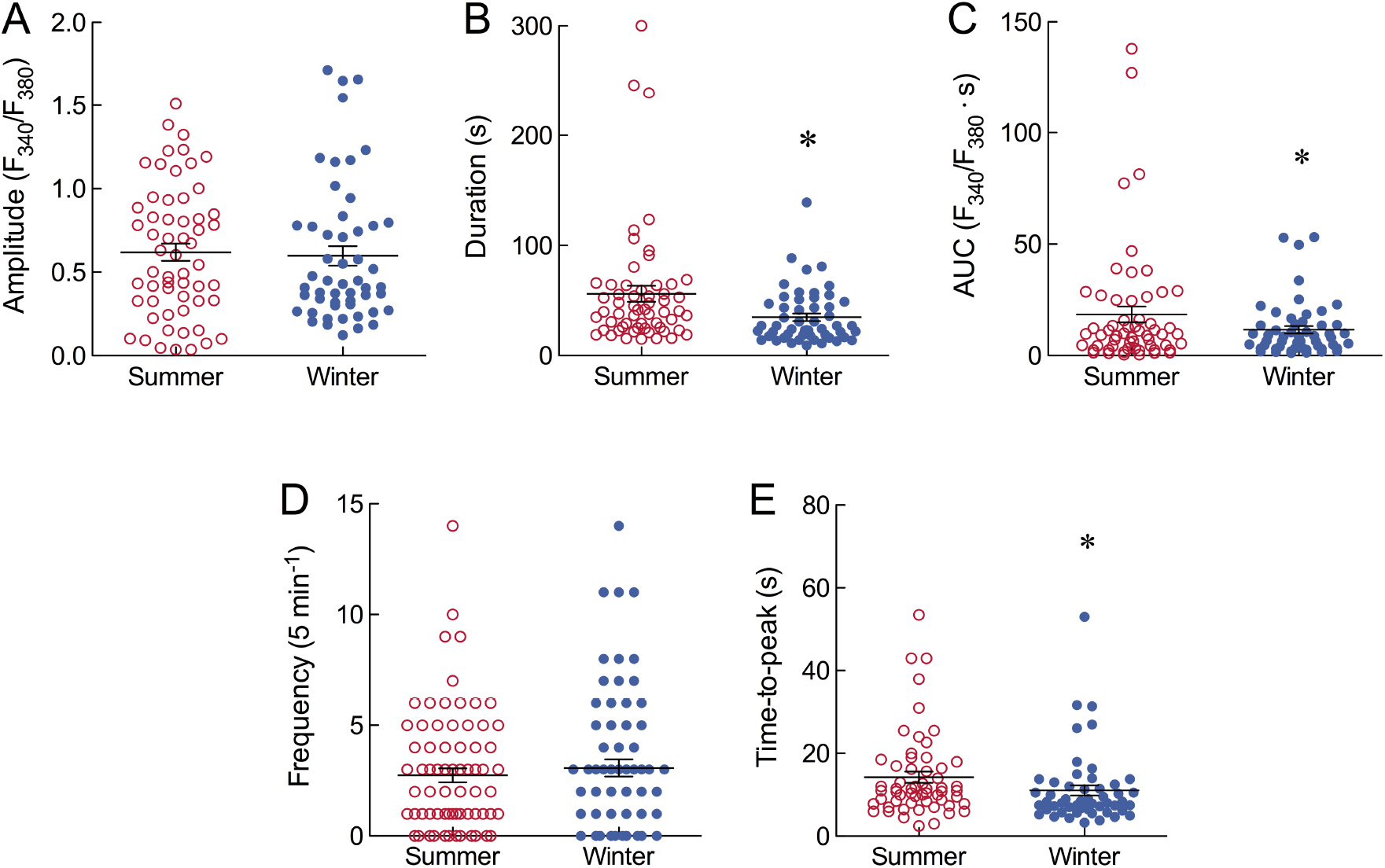
Summary data from parameter analysis of spontaneous Ca^2+^-based action potentials in cells isolated during the summer (open circles) and winter (closed circles). Shown are mean ± s.e.m. (A) Action potential amplitude was measured as the peak fluorescence ratio (F_340_/F_380_) from baseline: N = 59 cells in summer; 52 cells in winter. (B) Duration was measured in seconds (s): N = 59 cells in summer; 52 cells in winter. Asterisk indicates a significant difference (P = 0.0057, Student’s *t*-test). (C) Area under the curve (AUC) is an integral of F_340_/F_380_ with time: N = 59 cells in summer; 52 cells in winter. Asterisk indicates a significant difference (P = 0.049, Student’s *t*-test). (D) Frequency was recorded as events per 5- min sampling period: N = 76 cells in summer; 70 cells in winter. (E) Time-to-peak was the rise time from baseline to peak: N = 59 cells in summer; 52 cells in winter. Asterisk indicates a significant difference (P = 0.042, Student’s *t*-test).

Parameters of spontaneous action potentials, such as amplitude and frequency, were not significantly different between seasons (Fig. 7A,D; Student’s *t*-test). However, analysis of the duration of action potentials between seasons revealed that cells isolated during the summer had a significantly longer duration (56.1 ± 7.2 s) compared to those in winter (34.9 ± 3.4 s; P = 0.0057, Student’s *t*-test; Fig. 7B). In addition, AUC and time-to-peak were significantly greater in the summer compared to winter (P = 0.049 and P = 0.042, Student’s *t*-test; Fig. 7C,E), a trend that is consistent with longer duration action potential activity during the summer months.

## DISCUSSION

The present study demonstrates that seasonal changes in membrane structure and function in goldfish central neurons occur when animals are held under constant environmental conditions. We integrated techniques of mass spectrometry, atomic force microscopy and cell physiology to compare the chemical and biophysical properties of neurons isolated during the summer and winter months. Winter neurons displayed a greater proportion of unsaturated phosphatidylethanolamines and an increase in membrane stiffness. In addition, neurons isolated during the winter months exhibited reduced membrane excitability and shorter-duration Ca^2+^- based action potentials compared to summer neurons.

### Seasonal changes in membrane structure

Our analysis of membrane chemistry in whole retina using mass spectrometry revealed 153 species of phospholipids. Of these, only five species of PEs were found to change in concentration throughout the year, where PEs were highest in the winter compared to summer. In addition, the PEs that increased in concentration during the winter were unsaturated. The proportion of other phospholipids, such as PCs, did not exhibit seasonal change. Remodelling of the plasma membrane is an important aspect of thermal adaptation. PEs are disordering lipids, due to their conical shape, and decrease the degree of close packing of lipids within the bilayer compared to other phospholipids (Reynolds et al., 2014). It has previously been demonstrated that a reduction in temperature produces an elevated proportion of PEs relative to PCs, and an increase in unsaturated fatty acids (particularly long-chain polyunsaturated fatty acids) in cold-tolerant animals (Hazel, 1988; Pruitt, 1988; Hazel, 1995). Such changes are important for HVA to maintain constant viscosity or fluidity within the membrane as temperatures change. To our knowledge, this is the first report of the occurrence of HVA in vertebrate neurons in the absence of external cues, such as would be experienced during acclimatization to lower temperatures.

The seasonal changes in PEs that we observed in the goldfish retina represent only 3% of total phospholipid species, but suggest potentially important chemical modifications in the bilayer of retinal neurons that may occur before, or in parallel with, temperature-induced changes. Because HVA induced by temperature acclimatization may take weeks to occur (Cossins et al., 1977; Sellner and Hazel, 1982), a mechanism of temperature-independent changes in PEs may ensure that structural modifications in the membrane bilayer, which are critical for HVA, begin before environmental temperatures begin to change. This may provide cold-tolerant animals, such as goldfish, with an additional advantage for surviving cold temperatures by reducing the time required for HVA.

We found that winter neurons displayed an increase in membrane stiffness, as determined by nano-indentation by AFM and subsequent measurement of Young’s modulus. The mechanics of the plasma membrane and cortex, particularly actin filaments, are physically linked and so measurement of Young’s modulus will include local elastic responses of both membrane and cytoskeleton (Haase and Pelling, 2015). However, local measurements of Young’s modulus were taken over the soma, away from higher concentrations of actin filaments that were found primarily in horizontal cell dendrites. Although actin is present throughout somata and dendrites, a higher concentration of dendritic actin was previously described in horizontal cells of white bass (*Roccus chrysops*; Vaughan and Lasater, 1990). An increase in stiffness within the membrane bilayer correlates well with our observation of increased concentrations of PEs during the winter without changes in PC concentration. Moreover, an increase in the PE:PC ratio was recently shown to increase bilayer rigidity in liposomes, thereby maintaining membrane fluidity (Dawaliby et al., 2016). Al-Rekabi and Contera (2018) showed that, at high cholesterol concentrations, model lipid bilayers had a higher Young’s modulus and an increased membrane stiffness. Sterols can regulate membrane fluidity by interfering with acyl chain packing and reducing transitions to the solid state, but they can also increase rigidity of fluid membranes by limiting flexibility of nearby unsaturated acyl chains (Holthuis and Menon, 2014). The location of the nucleus within the cytosol may also impact measurement of Young’s modulus in whole-cell preparations (Haase and Pelling, 2015). In the present study, however, we performed nano- indentation experiments on the membrane over large central regions of the soma, away from the nucleus; and in control experiments, where we compared the Young’s modulus between on- and off-nucleus, we found no differences in our results.

The seasonal changes in membrane chemistry and stiffness reported in the present study are important because they may have a corresponding impact on the function of integral membrane proteins, such as ion channels. Membrane lipids modulate ion channels in one of two ways: through non-specific interactions—such as by changes in membrane curvature, thickness or fluidity—or through specific binding of select lipids to protein domains. Lipid-protein interactions may therefore lead to significant effects on ion channel activity and membrane excitability (Tillman and Cascio, 2003; Poveda et al., 2014; Rusinova et al., 2014). A change in the proportion of phospholipids with saturated versus unsaturated acyl chains, as occurs during HVA, may also impact the function of membrane proteins. Polyunsaturated fatty acids can modify gating and activation of voltage-gated ion channels (Moreno et al., 2012). In vertebrates, the degree of polyunsaturation in membrane phospholipids is also correlated with Na^+^/K^+^- ATPase and Ca^2+^-ATPase activity and maintenance of ionic gradients (Cossins et al., 1981; Hulbert and Else, 1999, 2005; Arnold et al., 2015). Importantly, naturally-occurring seasonal acclimatization, or laboratory-based thermal acclimation, have been shown to affect activity of ion channels in excitable membranes of fish (Hassinen et al., 2007, 2008; Rohmann et al., 2013; Kubly and Stecyk, 2015; Vornanen, 2016; Filatova et al., 2019), although these have not yet been linked specifically with structural changes in the phospholipid bilayer.

### Seasonal changes in membrane physiology

We found that neurons isolated during the winter had a reduced membrane excitability compared to summer neurons. We measured the parameters, *V*_rev_ and *R*_in_, as indicators of membrane excitability. *V*_rev_ is the voltage at which there is no net current measured across the membrane and provides an estimate of the resting membrane potential, which is not measured directly in voltage-clamp experiments. A shift in *V*_rev_ in the hyperpolarizing (i.e. leftward) direction on an *I-V* curve would represent a decrease in membrane excitability, as the resting potential moves further away from the activation threshold of voltage-dependent ion channels and generation of an action potential becomes less likely. *R*_in_ is inversely proportional to membrane conductance and is a convenient measure of ion channel activity. A membrane with a greater number of open channels would have a lower *R*_in_ and, according to Ohm’s Law, would be less excitable because a given inward current would have only a small depolarizing effect on the membrane potential. In the CNS and other excitable tissues, *R*_in_ and membrane excitability are controlled by K^+^ channels that are open at rest, such as the inwardly rectifying K^+^ (Kir) channels and leak K^+^ channels (Nichols and Lopatin, 1997; Reimann and Ashcroft, 1999; Goldstein et al., 2001; Lesage, 2003; Hibino et al., 2010).

Winter neurons had a *V*_rev_ of –80 mV, which occurred at a segment of the *I-V* curve that displayed a positive slope conductance as it crossed the voltage axis. This indicates a region of high membrane stability, where a shift in membrane potential in either direction would be expected to elicit an opposing current that would return the membrane potential to this resting state (Shingai and Christensen, 1986). Moreover, because *V*_rev_ was approximately equal to the calculated equilibrium potential for K^+^ in our experiments, it was evident that ionic permeability of membranes in winter was due primarily to K^+^ channels. Both voltage-gated and inwardly-rectifying K^+^ channels are present in teleost horizontal cells, although only Kir is substantially activated at resting potentials (Tachibana, 1983; Lasater, 1986; Jonz and Barnes, 2007). By contrast, summer neurons displayed inward (i.e. depolarizing) current at –80 mV and had correspondingly depolarized values of *V*_rev_. This suggests that neurons in summer maintained a relatively right-shifted resting membrane potential due to a relatively greater influence of other ion conductances on *V*_rev_.

In the retina, horizontal cells receive glutamatergic input from pre-synaptic photoreceptors. As a consequence, horizontal cells maintain a relatively depolarized resting membrane potential *in vivo* broadly estimated to be in the range of –35 mV to –20 mV, and hyperpolarize when photoreceptors are exposed to flashes of light (Shingai and Christensen, 1986; Yang et al., 1988; Thoreson and Mangel, 2012; Sun et al., 2017). Our experiments were performed on isolated cells, where resting membrane potentials would be relatively hyperpolarized in the absence of glutamate. Our results therefore indicate that seasonal modulation of membrane excitability must be an intrinsic property of these neurons. To address the issue of whether horizontal cells might respond differently to excitatory inputs in winter versus summer, we measured *R*_in_ over a range of voltages that correspond to resting potentials in the intact retina. We found that summer neurons had a higher *R*_in_ compared to those isolated during the winter. This suggests a higher level of membrane excitability in summer, as resting potentials would be closer to the activation potential of voltage-dependent Ca^2+^ channels (i.e. above –45 mV; Tachibana, 1983; Lasater, 1986; Jonz and Barnes, 2007).

Ca^2+^-based action potentials in horizontal cells have been described in a number of species, and are present in isolated cells, tissue slices and intact retina (Johnston and Lam 1981; Tachibana, 1981; Shingai and Christensen, 1983; Lasater et al. 1984; Murakami and Takahashi, 1987; Dixon et al., 1993; Takahashi et al. 1993; Kreitzer et al., 2012; Country et al., 2019, 2021). Although a specific role for action potentials in horizontal cells has not been firmly established, they may be involved in Ca^2+^-dependent regulation of pH (Kreitzer et al., 2012), they may be important for light-to-dark transitions in membrane potential, or play a role in feedback inhibition of pre-synaptic photoreceptors (Country et al., 2019). When monitoring intracellular Ca^2+^ dynamics, we found that spontaneous Ca^2+^-based action potentials in summer neurons were of longer duration and greater area under the curve (an integrated measure of intracellular Ca^2+^ activity over time). The increase in Ca^2+^ activity is consistent with a depolarized resting state and higher membrane excitability in summer neurons, where activation of voltage-dependent Ca^2+^ channels is more likely. We previously showed that increasing membrane excitability by injection of current to depolarize the membrane potential increased action potential activity in isolated horizontal cells (Country et al., 2019). As predicted by seasonal measurements of *V*_rev_, our Ca^2+^-imaging data therefore point to maintenance of a more depolarized resting membrane potential in summer neurons.

Kir is important in central neurons for setting resting membrane potential and controlling membrane excitability (Nichols and Lopatin, 1997; Reimann and Ashcroft, 1999; Hibino et al., 2010), and in horizontal cells Kir is also involved in accelerating the hyperpolarizing response to light (Dong and Werblin, 1995). In spiny neurons of the striatum, Kir hyperpolarizes the membrane potential to a “down state” and reduces *R*_in_, thereby producing a shunting effect that makes the cell relatively insensitive to small synaptic inputs (Nisenbaum and Wilson, 1995; John and Manchanda, 2011; Wilson, 2008). An analogous mechanism of shunting inhibition, though involving Cl^-^ conductance through GABA receptors, underlies suppression of action potentials in cortical neurons of anoxia-tolerant turtles (Pamenter et al., 2011). Moreover, a hyperpolarizing shift in *V*_rev_ by Kir was recently proposed to contribute to the temperature-dependent loss of electrical excitation in cardiac myocytes in fish, which may play a role in seasonal acclimatization (Vornanen et al., 2014; Vornanen, 2016; Badr et al., 2018). Kir channel expression has also been shown to be regulated by thermal acclimation (Hassinen et al., 2007, 2008; Vornanen, 2016). In addition, an increase in Kir current density and shortened action potential was reported in cardiac myocytes in the European shorthorn sculpin (*Myoxocephalus scorpio*) caught during the winter (Filatova et al., 2019).

We propose that, in a similar manner, Kir may modulate *R*_in_ and membrane excitability in goldfish horizontal cells during the winter to reduce action potential duration or depolarizing inputs from photoreceptors, thereby lowering energy requirements. Regulation of ionic gradients, such as following an action potential, consumes approximately 50% of neuronal ATP (Bickler and Buck, 2007; Howarth et al., 2012). Moreover, the unstimulated retina generally has high energy requirements due to photoreceptor dark current and high Na^+^/K^+^-ATPase activity (Ames, 1992; Country, 2017). Horizontal cells receive tonic stimulation from photoreceptors yet actively maintain transmembrane ionic gradients, regulate intracellular Ca^2+^ homeostasis and (in fish) support loading of synaptic vesicles with γ-aminobutyric acid (GABA) for Ca^2+^-dependent neurosecretion (Thoreson and Mangel, 2012; Country and Jonz, 2017)—all of which are mechanisms that consume energy in the form of ATP. Reduced activity in horizontal cells during the winter may, therefore, lower energy consumption in cold- and anoxia-tolerant animals, such as goldfish. In the congeneric crucian carp (*C. carassius*), exposure to anoxia reduced electroretinogram (ERG) activity by 90% (Johansson et al., 1997). A reduction in ERG activity by anoxia was also reported in other anoxia-tolerant vertebrates (Stensløkken et al., 2008), and by low temperature in hibernating ground squirrels (Zhang et al., 2020), suggesting that an overall reduction in retinal activity may be an important adaptation for survival in vertebrates that are tolerant to low oxygen or temperatures. Further investigation is needed, however, to determine whether the seasonal changes observed in the present study are confined to *Carassius* spp. and other vertebrates adapted to environmental changes, and whether they may be present in other neurons of the retina or CNS.

### What might be regulating seasonal changes in neuronal membranes?

In the present study, goldfish were imported at mixed age and from a natural photoperiod, but were maintained in our animal facilities at constant photoperiod, temperature and oxygen availability. Although we have not yet explored what might be directly regulating the observed seasonal patterns in membrane structure and excitability, we suspect that biological rhythms associated with seasonal or annual cycles were entrained before fish were imported into our holding facilities. Seasonal changes in photoperiod and temperature regulate many important physiological processes, such as growth, feeding, metabolism and reproduction (Peter and Crim, 1979; Zhang et al., 2009; Cowan et al., 2017; Nakane and Yoshimura, 2019). In goldfish, reproductive ability is maintained in captivity, where gene expression patterns in the brain, including genes involved in neurotransmission, change between May and August, which corresponds to a transition from prespawning to sexual regression (Zhang et al., 2009).

Furthermore, there is evidence that reproductive transitions are entrained in fish and continue for multiple cycles without the need for environmental cues. Duston and Bromage (1991) found that 2-year-old rainbow trout (*Oncorhynchus mykiss*) subsequently maintained in captivity at constant photoperiod and temperature continued to display seasonal rhythms of gonadal maturation and ovulation for up to 3 years. Further investigation is required to establish whether seasonal changes in gene expression that modify phospholipid composition and membrane excitability are entrained in neurons of vertebrates adapted to extreme environments, and to potentially determine the longevity of these cyclical patterns in other, potentially longer-living species. In addition, it will be of interest to explore whether regulation of specific phospholipid-protein signalling mechanisms might impact seasonal changes in membrane excitability. For example, the membrane phospholipid, phosphoinositol 4,5-bisphosphate (PIP_2_), is essential for maintaining activity of most Kir channels (reviewed by Hibino et al., 2010), and an increase in PIP_2_ activity was associated with the hyperpolarizing response to light in horizonal cells of *Xenopus laevis* (Anderson and Hollyfield, 1981, 1984). Interestingly, seasonal changes in the regulation of ion channels by PIP_2_ in dorsal root ganglion cells modulates thermal sensitivity in mice during adaptation to changing ambient temperatures (Fujita et al., 2013; Brenner et al., 2014). Whether direct interactions between PIP_2_, or other membrane phospholipids, and ion channels are involved in controlling seasonal changes in membrane excitability in the goldfish retina will be an important avenue for further research. Future studies focused on identifying the role of seasonal patterns controlling membrane excitability in central neurons will provide a better understanding of the evolutionary adaptations that allow some organisms to survive extreme environmental change.

## Acknowledgements

The authors thank Cameron Jonz for colour modifications to Fig. 4B; and Dr. Steven Barnes for access to animals and equipment at the Department of Physiology and Biophysics, Dalhousie University.

## Author contributions

Conceptualization: M.G.J., J.C.S., A.E.P.; Methodology: all authors; Validation: M.G.J., M.W.C., K.H., K.B., B.F.N.C., J.C.S.; Formal analysis: M.G.J., M.W.C., K.H., K.B., C.R.C., J.A.B., J.C.S.; Investigation: all authors.; Resources: M.G.J., J.C.S., A.E.P; Data curation: M.G.J., M.W.C., K.H., K.B., C.R.C., J.A.B., J.C.S.; Writing - original draft: M.G.J., M.W.C., K.H., K.B., J.C.S.; Writing - review & editing: all authors; Supervision: M.G.J., J.C.S., A.E.P.; Project administration: M.G.J.; Funding acquisition: M.G.J., J.C.S., A.E.P.

## Data availability

All data associated with this work are presented in the manuscript.

## References

Akopian, A., McReynolds, J. and Weiler, R. (1991). Short-term potentiation of off-responses in turtle horizontal cells. Brain Res. 546, 132–138

Al-Rekabi, Z. and Contera, S. (2018). Multifrequency AFM reveals lipid membrane mechanical properties and the effect of cholesterol in modulating viscoelasticity. Proc. Natl. Acad. Sci. U S A. 115, 2658–2663. doi:10.1073/pnas.1719065115

Ames, A., Li, Y.Y., Heher, E.C. and Kimble, C.R. (1992). Energy metabolism of rabbit retina as related to function: high cost of Na^+^ transport. J. Neurosci. 12, 840–853

Andersen, O.S. and Koeppe, R.E. (2007). Bilayer thickness and membrane protein function: an energetic perspective. Annu. Rev. Biophys. Biomol. Struct. 36, 107–130

Anderson, R.E. and Hollyfield, J.G. (1981). Light stimulates the incorporation of inositol into phosphatidylinositol in the retina. Biochim. Biophys Acta. 665, 619–622

Anderson, R.E. and Hollyfield, J.G. (1984). Inositol incorporation into phosphoinositides in retinal horizontal cells of *Xenopus laevis*: enhancement by acetylcholine, inhibition by glycine. J Cell Biol. 99, 686–691

Arnold, W., Giroud, S., Valencak, T.G. and Ruf, T. (2015). Ecophysiology of omega fatty acids: a lid for every jar. Physiology (Bethesda). 30, 232–240

Badr, A., Korajoki, H., Abu-Amra, E.S., El-Sayed, M.F. and Vornanen, M. (2018). Effects of seasonal acclimatization on thermal tolerance of inward currents in roach (*Rutilus rutilus*) cardiac myocytes. J. Comp. Physiol. B. 188, 255–269. doi: 10.1007/s00360-017-1126-1

Benjamini, Y. and Hochberg, Y. (1995). Controlling the false discovery rate: a practical and powerful approach to multiple testing. J. R. Statist. Soc. B. 57, 289–300

Bickler, P.E. and Buck, L.T. (2007). Hypoxia tolerance in reptiles, amphibians, and fishes: life with variable oxygen availability. Annu. Rev. Physiol. 69, 145–170

Bligh, E.G. and Dyer, W.J. (1959). A rapid method of total lipid extraction and purification. Can. J. Biochem. Physiol. 37, 911–917

Brenner, D.S., Golden, J.P., Vogt, S.K., Dhaka, A., Story, G.M. and Gereau, R.W. IV. (2014). A dynamic set point for thermal adaptation requires phospholipase C-mediated regulation of TRPM8 *in vivo*. Pain. 155, 2124–2133

Canez, C.R., Shields, S.W., Bugno, M., Wasslen, K.V., Weinert, H.P., Willmore, W.G., Manthorpe, J.M. and Smith, J.C. (2016). Trimethylation enhancement using (13)C- diazomethane ((13)C-TrEnDi): increased sensitivity and selectivity of phosphatidylethanolamine, phosphatidylcholine, and phosphatidylserine lipids derived from complex biological samples. Anal. Chem. 88, 6996–7004

Carl, P., Kwok, C.H., Manderson, G., Speicher, D.W. and Discher, D.E. 2001. Forced unfolding modulated by disulfide bonds in the Igdomains of a cell adhesion molecule. Proc. Natl. Acad. Sci. USA. 98, 1565–1570

Chambers, M.C., Maclean, B., Burke, R., Amodei, D., Ruderman, D.L., Neumann, S., Gatto, L., Fischer, B., Pratt, B., Egertson, J. et al. (2012). A cross-platform toolkit for mass spectrometry and proteomics. Nat .Biotechnol. 30, 918–920

Cossins, A.R., Bowler, K. and Prosser, C.L. (1981). Homeovisous adaptation and its effect upon membrane-bound proteins. J. Therm. Biol. 6, 183–187

Cossins, A.R., Friedlander, M.J. and Prosser, C.L. (1977). Correlations between behavioral temperature adaptations of goldfish and the viscosity and fatty acid composition of their synaptic membranes. J. Comp. Physiol. 120, 109–121

Country, M.W. (2017). Retinal metabolism: a comparative look at energetics in the retina. Brain Res. 1672, 50–57

Country, M.W. and Jonz, M.G. (2017). Calcium dynamics and regulation in horizontal cells of the vertebrate retina: lessons from teleosts. J. Neurophysiol. 117, 523–536

Country, M.W., Campbell, B.F.N. and Jonz, M.G. (2019). Spontaneous action potentials in retinal horizontal cells of goldfish (*Carassius auratus*) are dependent upon L-type Ca^2+^ channels and ryanodine receptors. J. Neurophysiol. 122, 2284–2293

Country, M.W., Htite, E.D., Samson, I.A. and Jonz, M.G. (2021). Retinal horizontal cells of goldfish (*Carassius auratus*) display subtype-specific differences in spontaneous action potentials *in situ*. J. Comp. Neurol. 529, 1756–1767. doi: 10.1002/cne.25054

Cowan, M., Azpeleta, C. and López-Olmeda, J.F. (2017). Rhythms in the endocrine system of fish: a review. J. Comp. Physiol. B. 187, 1057–1089

Cunningham, J.R. and Neal, M.J. (1985). Effect of excitatory amino acids on gamma-aminobutyric acid release from frog horizontal cells. J. Physiol. 362, 51–67

Dawaliby, R., Trubbia, C., Delporte, C., Noyon, C., Ruysschaert, J.M., Van Antwerpen, P. and Govaerts, C. (2016). Phosphatidylethanolamine is a key regulator of membrane fluidity in eukaryotic cells. J. Biol. Chem. 291, 3658–3667

DeVries, S.H. and Schwartz, E.A. (1992). Hemi-gap-junction channels in solitary horizontal cells of the catfish retina. J. Physiol. 445, 201–230

Dixon, D.B., Takahashi, K. and Copenhagen, D.R. (1993). L-Glutamate suppresses HVA calcium current in catfish horizontal cells by raising intracellular proton concentration. Neuron. 11, 267–277

Dong, C.J. and Werblin, F.S. (1995). Inwardly rectifying potassium conductance can accelerate the hyperpolarizing response in retinal horizontal cells. J. Neurophysiol. 74, 2258–2265

Dowling, J.E. (1987). The Retina: An Approachable Part of the Brain. Cambridge, MA: Harvard University Press

Dowling, J.E., Pak, M.W. and Lasater, E. (1985). White perch horizontal cells in culture: methods, morphology and process growth. Brain Res. 360: 331–338

Duston, J. and Bromage, N. (1991). Circannual rhythms of gonadal maturation in female rainbow trout (*Oncorhynchus mykiss*). J. Biol. Rhythms. 6, 49–53

Ernst, R., Ejsing, C.S. and Antonny, B. Homeoviscous adaptation and the regulation of membrane lipids. J. Mol. Biol. 428, 4776–4791

Fhaner, C.J., Liu, S., Ji, H., Simpson, R.J. and Reid, G.E. (2012). Comprehensive lipidome profiling of isogenic primary and metastatic colon adenocarcinoma cell lines. Anal. Chem. 84, 8917–8926

Filatova, T.S., Abramochkin, D.V. and Shiels, H.A. (2019). Thermal acclimation and seasonal acclimatization: a comparative study of cardiac response to prolonged temperature change in shorthorn sculpin. J. Exp. Biol. 222, jeb202242. doi: 10.1242/jeb.202242

Fried, S.D.E, Lewis, J.W., Szundi, I., Martinez-Mayorga, K., Mahalingam, M., Vogel, R., Kliger, D.S. and Brown, M.F. (2021). Membrane curvature revisited-the archetype of rhodopsin studied by time-resolved electronic spectroscopy. Biophys. J. 120, 440–452

Fujita, F., Uchida, K., Takaishi, M., Sokabe, T. and Tominaga, M. (2013). Ambient temperature affects the temperature threshold for TRPM8 activation through interaction of phosphatidylinositol 4,5-bisphosphate. J. Neurosci. 33, 6154–6159

Goldstein, S.A., Bockenhauer, D., O’Kelly, I. and Zilberberg, N. (2001). Potassium leak channels and the KCNK family of two-P-domain subunits. Nat. Rev. Neurosci. 2, 175–184

Haase, K. and Pelling, A.E. (2013). Resiliency of the plasma membrane and actin cortex to large-scale deformation. Cytoskeleton (Hoboken). 70, 494–514

Haase, K. and Pelling, A.E. (2015). Investigating cell mechanics with atomic force microscopy. J. R. Soc. Interface. 12(104):20140970. doi:10.1098/rsif.2014.0970

Hassinen, M., Paajanen, V. and Vornanen, M. (2008). A novel inwardly rectifying K^+^ channel, Kir2.5, is upregulated under chronic cold stress in fish cardiac myocytes. J. Exp. Biol. 211, 2162–2171

Hassinen, M., Paajanen, V., Haverinen, J., Eronen, H. and Vornanen, M. (2007). Cloning and expression of cardiac Kir2.1 and Kir2.2 channels in thermally acclimated rainbow trout. Am. J. Physiol. Regul. Integr. Comp. Physiol. 292, R2328–2339

Hazel, J.R. (1988). Homeoviscous adaptation in animal cell membranes. In Advances in Membrane Fluidity—Physiological Regulation of Membrane Fluidity (ed. R.C. Aloia, C.C. Curtain and L.M. Gordon), 6:149–88. New York: Liss

Hazel, J.R. (1995). Thermal adaptation in biological membranes: is homeoviscous adaptation the explanation? Annu. Rev. Physiol. 57, 19–42

Hazel, J.R. and Williams, E.E. (1990). The role of alterations in membrane lipid composition in enabling physiological adaptation of organisms to their physical environment. Prog. Lipid. Res. 29, 167–227

Herbert, C.V. and Jackson, D.C. (1985). Temperature effects on the responses to prolonged submergence in the turtle. *Chrysemys picta bellii*. Physiol. Zool. 58, 670–681

Hibino, H., Inanobe, A., Furutani, K., Murakami, S., Findlay, I. and Kurachi, Y. (2010). Inwardly rectifying potassium channels: their structure, function, and physiological roles. Physiol. Rev. 90, 291–366

Hille, B. (2001). Ion Channels of Excitable Membranes, third ed. Sunderland, MA: Sinauer.

Hochachka, P.W., Buck, L.T., Doll, C.J. and Land, S.C. (1996). Unifying theory of hypoxia tolerance: molecular/metabolic defense and rescue mechanisms for surviving oxygen lack. Proc. Natl. Acad. Sci. USA. 93, 9493–9498

Holthuis, J.C. and Menon, A.K. (2014). Lipid landscapes and pipelines in membrane homeostasis. Nature. 510, 48–57

Howarth, C., Gleeson, P. and Attwell, D. (2012). Updated energy budgets for neural computation in the neocortex and cerebellum. J. Cereb. Blood Flow Metab. 32, 1222–1232

Hulbert, A.J. and Else, P.L. (1999). Membranes as possible pacemakers of metabolism. J. Theor. Biol. 199, 257–274

Hulbert, A.J. and Else, P.L. (2005). Membranes and the setting of energy demand. J. Exp. Biol. 208, 1593–1599

Jackson, D.C. and Ultsch, G.R. (2010). Physiology of hibernation under the ice by turtles and frogs. J. Exp. Zool. 313A, 311-327

Jackson, D.C., Herbert, C.V. and Ultsch, G.R. (1984). The comparative physiology of diving in North American freshwater turtles. II. Plasma ion balance during prolonged anoxia. Physiol. Zool. 57, 632–640

Johansson, D., Nilsson, G.E. and Døving, K.B. (1997). Anoxic depression of light-evoked potentials in retina and optic tectum of crucian carp. Neurosci. Lett. 237, 73–76

John, J. and Manchanda, R. (2011). Modulation of synaptic potentials and cell excitability by dendritic KIR and KAs channels in nucleus accumbens medium spiny neurons: a computational study. J. Biosci. 36, 309–328

Johnston, D. and Lam, D.M. (1981). Regenerative and passive membrane properties of isolated horizontal cells from a teleost retina. Nature. 292, 451–454

Jonz, M.G. and Barnes, S. (2007). Proton modulation of ion channels in isolated horizontal cells of the goldfish retina. J. Physiol. 581, 529–541

Kind, T., Liu, K.H., Lee, D.Y., DeFelice, B., Meissen, J.K. and Fiehn, O. (2013). LipidBlast *in silico* tandem mass spectrometry database for lipid identification. Nat. Methods. 10, 755–758

Kreitzer, M.A., Jacoby, J., Naylor, E., Baker, A., Grable, T., Tran, E., Booth, S.E., Qian, H. and Malchow, R.P. (2012). Distinctive patterns of alterations in proton efflux from goldfish retinal horizontal cells monitored with self-referencing H⁺-selective electrodes. Eur. J. Neurosci. 36, 3040–3050

Kubly, K.L. and Stecyk, J.A. (2015). Temperature-dependence of L-type Ca^2+^ current in ventricular cardiomyocytes of the Alaska blackfish (*Dallia pectoralis*). J. Comp. Physiol. B. 185, 845–858

Lasater, E.M. (1986). Ionic currents of cultured horizontal cells isolated from white perch retina. J. Neurophysiol. 55, 499–513

Lasater, E.M., Dowling, J.E. and Ripps, H. (1984). Pharmacological properties of isolated horizontal and bipolar cells from the skate retina. J. Neurosci. 4, 1966–1975

Lee, A.G. (2004). How lipids affect the activities of integral membrane proteins. Biochim. Biophys. Acta. 1666, 62–87

Lesage F. (2003). Pharmacology of neuronal background potassium channels. Neuropharmacology. 44, 1–7

Lutz, P.L. and Nilsson, G.E. (2004). Vertebrate brains at the pilot light. Respir. Physiol. Neurobiol. 141, 285–296

Malchow, R.P., Qian, H.H., Ripps, H. and Dowling, J.E. (1990). Structural and functional properties of two types of horizontal cell in the skate retina. J. Gen. Physiol. 95, 177–198

Moreno, C., Macias, A., Prieto, A., De La Cruz, A. and Valenzuela, C. (2012). Polyunsaturated fatty acids modify the gating of kv channels. Front. Pharmacol. 3:163. doi: 10.3389/fphar.2012.00163

Murakami, M. and Takahashi, K. (1987). Calcium action potential and its use for measurement of reversal potentials of horizontal cell responses in carp retina. J. Physiol. 386, 165–180

Nakane, Y. and Yoshimura, T. (2019). Photoperiodic regulation of reproduction in vertebrates. Annu. Rev. Anim. Biosci. 7, 173–194

Nichols, C.G. and Lopatin, A.N. (1997). Inward rectifier potassium channels. Annu. Rev. Physiol. 59, 171–191

Nisenbaum, E.S. and Wilson, C.J. (1995). Potassium currents responsible for inward and outward rectification in rat neostriatal spiny projection neurons. J. Neurosci. 15, 4449–4463

Pamenter, M.E., Shin, D.S., Cooray, M. and Buck, L.T. (2008). Mitochondrial ATP-sensitive K^+^ channels regulate NMDAR activity in the cortex of the anoxic western painted turtle. J. Physiol. 586, 1043–1058

Pamenter, M.E., Hogg, D.W., Ormond, J., Shin, D.S., Woodin, M.A. and Buck, L.T. (2011). Endogenous GABA_A_ and GABA_B_ receptor-mediated electrical suppression is critical to neuronal anoxia tolerance. Proc. Natl. Acad. Sci. USA. 108, 11274–11279

Peter, R.E. and Crim, L.W. (1979). Reproductive endocrinology of fishes: gonadal cycles and gonadotropin in teleosts. Annu. Rev. Physiol. 41, 323–335

Pluskal, T., Castillo, S., Villar-Briones, A., Oresic, M. (2010). MZmine 2: modular framework for processing, visualizing, and analyzing mass spectrometry-based molecular profile data. BMC Bioinformatics. 23;11:395. doi: 10.1186/1471-2105-11-395

Poveda, J.A., Giudici, A.M., Renart, M.L., Molina, M.L., Montoya, E., Fernández-Carvajal, A., Fernández-Ballester, G., Encinar, J.A. and González-Ros, J.M. (2014). Lipid modulation of ion channels through specific binding sites. Biochim. Biophys. Acta. 1838, 1560–1567

Pruitt, N.L. (1988). Membrane lipid composition and overwintering strategy in thermally acclimated crayfish. Am. J. Physiol. 254, R870–876

Reimann, F. and Ashcroft, F.M. (1999). Inwardly rectifying potassium channels. Curr. Opin. Cell Biol. 11, 503–508

Reynolds, A.M., Lee, R.E. Jr and Costanzo, J.P. (2014). Membrane adaptation in phospholipids and cholesterol in the widely distributed, freeze-tolerant wood frog, *Rana sylvatica*. J. Comp. Physiol. B. 184, 371–383

Rodgers-Garlick, C.I., Hogg, D.W. and Buck, L.T. (2013). Oxygen-sensitive reduction in Ca^2+^-activated K^+^ channel open probability in turtle cerebrocortex. Neuroscience. 237, 243–254

Rohmann, K.N., Fergus, D.J. and Bass, A.H. (2013). Plasticity in ion channel expression underlies variation in hearing during reproductive cycles. Curr. Biol. 23, 678–683

Rusinova, R., Kim, D.M., Nimigean, C.M. and Andersen, O.S. (2014). Regulation of ion channel function by the host lipid bilayer examined by a stopped-flow spectrofluorometric assay. Biophys. J. 106, 1070–1078

Schneider, C.A., Rasband, W.S. and Eliceiri, K.W. (2012). NIH Image to ImageJ: 25 years of image analysis. Nat. Methods. 9, 671–675

Sellner, P.A. and Hazel, J.R. (1982). Time course of changes in fatty acid composition of gills and liver from rainbow trout (*Salmo gairdneri*) during thermal acclimation. J. Exp. Zool. 221, 159–168

Shingai, R. and Christensen, B.N. (1986). Excitable properties and voltage-sensitive ion conductances of horizontal cells isolated from catfish (*Ictalurus punctatus*) retina. J. Neurophysiol. 56, 32–49

Sinensky, M. (1974). Homeoviscous adaptation—a homeostatic process that regulates the viscosity of membrane lipids in *Escherichia coli*. Proc. Natl. Acad. Sci. USA. 71, 522–525. doi: 10.1073/pnas.71.2.522

Stensløkken, K.O., Milton, S.L., Lutz, P.L., Sundin, L., Renshaw, G.M., Stecyk, J.A. and Nilsson, G.E. (2008). Effect of anoxia on the electroretinogram of three anoxia-tolerant vertebrates. Comp. Biochem. Physiol. A Mol. Integr. Physiol. 150, 395–403

Sun, X., Hirano, A.A., Brecha, N.C. and Barnes, S. (2017). Calcium-activated BK_Ca_ channels govern dynamic membrane depolarizations of horizontal cells in rodent retina. J. Physiol. 595, 4449–4465

Tachibana, M. (1981). Membrane properties of solitary horizontal cells isolated from goldfish retina. J. Physiol. 321, 141–161

Tachibana, M. (1983). Ionic currents of solitary horizontal cells isolated from goldfish retina. J. Physiol. 345, 329–351

Takahashi, K., Dixon, D.B. and Copenhagen, D.R. (1993). Modulation of a sustained calcium current by intracellular pH in horizontal cells of fish retina. J. Gen. Physiol. 101, 695–714

Thoreson, W.B. and Mangel, S.C. (2012). Lateral interactions in the outer retina. Prog. Retin. Eye Res. 31, 407–441

Tillman, T.S. and Cascio, M. (2003). Effects of membrane lipids on ion channel structure and function. Cell Biochem. Biophys. 38, 161–190

Vance, J.E. and Tasseva, G. (2013). Formation and function of phosphatidylserine and phosphatidylethanolamine in mammalian cells. Biochim. Biophys. Acta. 1831, 543–554

van den Thillart, G., Kesbeke, F. and van Waarde, A. (1980) Anaerobic energy-metabolism of goldfish, *Carassius auratus* (L.) Influence of hypoxia and anoxia on phosphorylated compounds and glycogen. J. Comp. Physiol. B. 136, 45–52

Vaughan, D.K. and Lasater, E.M. (1990). Distribution of F-actin in bipolar and horizontal cells of bass retinas. Am. J. Physiol. 259, C205–214

Vornanen, M. (2016). The temperature dependence of electrical excitability in fish hearts. J. Exp. Biol. 219, 1941–1952. doi: 10.1242/jeb.128439

Vornanen, M., Haverinen, J. and Egginton, S. (2014). Acute heat tolerance of cardiac excitation in the brown trout (*Salmo trutta fario*). J. Exp. Biol. 217, 299–309

Wilson, C. (2008). Up and down states. Scholarpedia J. 3(6):1410. doi: 10.4249/scholarpedia.1410

Yang, X.L., Tornqvist, K. and Dowling, J.E. (1988). Modulation of cone horizontal cell activity in the teleost fish retina. I. Effects of prolonged darkness and background illumination on light responsiveness. J. Neurosci. 8, 2259–2268

Zhang, H., Sajdak, B.S., Merriman, D.K., McCall, M.A., Carroll, J. and Lipinski, D.M. (2020). Electroretinogram of the cone-dominant thirteen-lined ground squirrel during euthermia and hibernation in comparison with the rod-dominant brown Norway Rat. Invest. Ophthalmol. Vis. Sci. 61, 6. doi: 10.1167/iovs.61.6.6

Zhang, D., Xiong, H., Mennigen, J.A., Popesku, J.T., Marlatt, V.L., Martyniuk, C.J., Crump, K., Cossins, A.R., Xia, X. and Trudeau, V.L. (2009). Defining global neuroendocrine gene expression patterns associated with reproductive seasonality in fish. PLoS One. 4(6):e5816. doi: 10.1371/journal.pone.0005816

Zivkovic, G. and Buck, L.T. (2010). Mitochondrial ATP sensitive K^+^ channels decrease AMPAR currents in the anoxic turtle cortex. J. Neurophysiol. 104, 1913–1922

